# Epigenetic priming of neural progenitors by Notch enhances Sonic hedgehog signaling and establishes gliogenic competence

**DOI:** 10.1101/2025.01.20.633996

**Authors:** Luuli N. Tran, Ashwini Shinde, Kristen H. Schuster, Aiman Sabaawy, Emily Dale, Madalynn J. Welch, Trevor J. Isner, Sylvia A. Nunez, Fernando García-Moreno, Charles G. Sagerström, Bruce H. Appel, Santos J. Franco

## Abstract

The remarkable cell diversity of multicellular organisms relies on the ability of multipotent progenitor cells to generate distinct cell types at the right times and locations during embryogenesis. A key question is how progenitors establish competence to respond to the different environmental signals required to produce specific cell types at critical developmental timepoints. We addressed this in the mouse developing forebrain, where neural progenitor cells must switch from producing neurons to making oligodendrocytes in response to increased Sonic Hedgehog (SHH) signaling during late embryogenesis. We show that progenitor responses to SHH are regulated by Notch signaling, thus permitting proper timing of the neuron-oligodendrocyte switch. Notch activity epigenetically primes genes associated with the oligodendrocyte lineage and SHH pathway, enabling amplified transcriptional responses to endogenous SHH and robust oligodendrogenesis. These results reveal a critical role for Notch in facilitating progenitor competence states and influencing cell fate transitions at the epigenetic level.

## INTRODUCTION

Cell fate specification is a critical process by which multipotent progenitor cells acquire distinct identities with specialized functions. To achieve this, progenitors must respond appropriately to environmental signals to generate the right cell types at the proper times during embryogenesis. A prime example of this process is found in the developing central nervous system, where multiple different types of neurons and glial cells originate from a shared pool of neural progenitor cells in a spatially and temporally regulated manner. In the developing dorsal forebrain, neural progenitors first generate excitatory neurons before they start producing glial cells like astrocytes and oligodendrocytes, in a process known as the “neuron-glia switch”^1–3^. This switch provides an excellent model to investigate the mechanisms that govern the acquisition of diverse fates from a common pool of neural progenitors.

Proper timing of the neuron-glia switch relies on precisely coordinated interactions between cell-intrinsic factors, such as epigenetic state and gene expression profile, and extracellular signals that instruct progenitors to commence a gliogenic transcriptional program^1–3^. For example, during early embryonic development the morphogen signaling molecule Sonic hedgehog (SHH) is highly expressed in the ventral forebrain to pattern the dorsal-ventral axis, allowing for neurogenesis to occur in dorsal regions^4^. Studies from our lab showed that SHH signaling increases in the dorsal forebrain during later stages of embryonic development, initiating the transition from neurogenesis to the formation of oligodendrocyte lineage cells^5^. Importantly, we found that the beginning of oligodendrogenesis overlaps with the end of neurogenesis, so that only some progenitors form oligodendrocytes in response to SHH ligand, while others continue to produce neurons. This suggests that additional pathways are required to make progenitors competent to produce oligodendrocyte lineage cells upon SHH exposure.

Multiple studies have demonstrated that the intrinsic properties of neural progenitors change over developmental time, and thus their competence to respond to specific extracellular signals also changes^6–9^. Although neural progenitors maintain a core cellular identity, single cell transcriptomics studies suggest that progenitors are transcriptionally primed with a mixed cell identity before a specific neuronal subtype lineage is resolved^6, 8^. Additionally, the chromatin landscape provides another key regulatory step in imposing divergent cell fates. For instance, temporal changes in chromatin dynamics of neural progenitors determine whether they generate neurons or astrocytes in response to BMP signaling^7^. These studies point towards the intriguing possibility that neural progenitors may require a specific intrinsic molecular state that is competent to produce oligodendrocyte lineage cells in response to SHH signaling.

We previously demonstrated that the Notch signaling pathway both promotes a progenitor state and is required for the proper timing and scale of dorsal forebrain oligodendrogenesis^10^. Here, we show that Notch signaling is required for SHH-mediated oligodendrogenesis, providing us with a framework to interrogate the intrinsic progenitor state that is competent to respond to SHH and to specify the oligodendrocyte lineage. We used Notch gain-of-function mutants to profile the transcriptomes and chromatin landscapes of neural progenitors experiencing Notch pathway over-activation during the neuron-glia switch in the dorsal forebrain. We found that Notch activation reorganized the genome in favor of a more gliogenic and less neurogenic chromatin state, and promoted accessibility of chromatin nearby SHH pathway genes. Interestingly, at the transcriptional level higher Notch signaling initially promoted an undifferentiated progenitor identity that lacked expression of mRNAs known to drive cell fate specification and differentiation. However, a day later during the peak of the neuron-glia switch, Notch activation increased oligodendrogenesis. Finally, mRNA sequencing revealed upregulation of SHH pathway components, and *in vivo* reporter assays revealed that Notch activation amplified SHH transcriptional output. These data illustrate a model in which Notch signaling maintains an undifferentiated epigenetic and transcriptomic state in neural progenitors, while also establishing a chromatin landscape that inhibits neurogenesis and primes progenitors for a gliogenic program. Later, primed progenitors experience an enhanced response to SHH signaling and initiate robust oligodendrocyte production, thus contributing to the proper timing and scale of the neuron-glia switch.

## RESULTS

### Notch signaling regulates the production of oligodendrocyte lineage cells in response to SHH

We previously showed that the SHH and Notch signaling pathways are individually critical for proper oligodendrogenesis from dorsal forebrain progenitors during the neuron-glia switch^5, 10^. Given that Notch signaling can impact SHH signaling in other developmental contexts^11–16^, we hypothesized that Notch pathway activation controls oligodendrogenesis in part by regulating progenitor responses to SHH. To test this hypothesis, we first asked if Notch signaling is required for SHH-induced oligodendrogenesis using an *ex vivo* forebrain slice culture system (**Figure 1**). Dorsal forebrain progenitors begin generating the oligodendrocyte lineage during late embryogenesis, starting with the production of OLIG2+ glial progenitor cells, some of which then express PDGFRA as they transition to oligodendrocyte precursor cells (OPCs)^5, 17^ (**Figure 1A**). We cut forebrain slices from mouse embryos right before the neuron-glia switch at embryonic day (E) 15.5 and cultured them for 2 days to analyze oligodendrogenesis by staining for OLIG2 and PDGFRA (**Figure 1B**). Following Notch receptor activation, γ-secretases cleave the Notch intracellular domain (NICD), which forms a transcriptional complex with the cofactor RBPJ to activate target gene expression^18^. We blocked Notch signaling in slice cultures by adding the γ-secretase inhibitor DAPT and stimulated the SHH pathway by adding recombinant SHH ligand (**Figure 1B**). We previously showed that DAPT treatment severely reduces the number of OLIG2+ cells in the pallium^10^. In contrast, we found here that SHH treatment significantly increased OLIG2+ cells by twofold, consistent with its role in promoting dorsal oligodendrogenesis. However, this SHH-induced increase in OLIG2+ cells was completely blocked when slices were treated with both SHH and DAPT together (**Figures 1C and 1D**). Conversely, we found the opposite when Notch signaling was activated. To activate the Notch pathway specifically in the dorsal lineage, we crossed *Emx1-Cre* mice^19^ to the *R26-Lox-Stop-Lox-NICD* line^20^. The resulting mice, hereafter referred to as NICD, overexpress NICD in *Emx1*+ dorsal forebrain progenitors^21^. We found that adding SHH to slices from NICD brains resulted in an even greater increase in OLIG2+ cells compared to SHH alone (**Figures 1C and 1D**). Together, these data demonstrate that SHH-mediated oligodendrogenesis in the dorsal forebrain can be modulated by Notch signaling.

**Figure 1.**
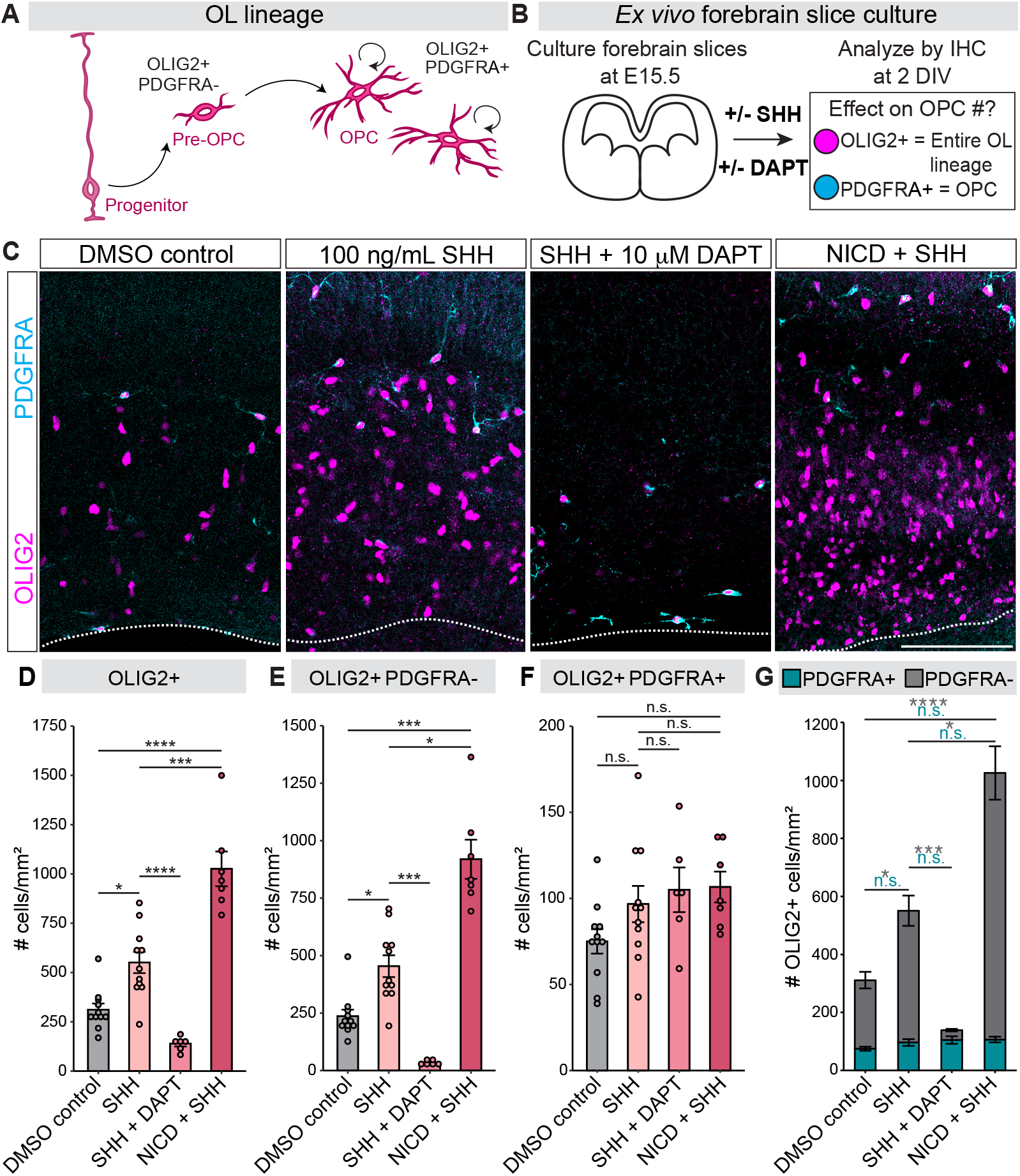
Notch pathway inhibition blocks SHH-induced oligodendrogenesis in forebrain slice cultures. (A) Schematic of OL lineage progression. Pre-OPCs derived from dorsal forebrain progenitors express OLIG2 and transition into OPCs once they start expressing PDGFRA. OPCs proliferate and populate the dorsal pallium. OL, oligodendrocyte; OPC, oligodendrocyte precursor cell. (B) Schematic of forebrain slice culture approach. Brains of E15.5 wildtype or NICD mouse embryos were sectioned and cultured with or without DAPT and SHH. At 2 DIV, forebrain slices were fixed and stained for OLIG2 and PDGFRA to identify oligodendrocyte lineage cells. DIV = days *in vitro*; IHC = immunohistochemistry. (C) Representative images of OLIG2+ and OLIG2+ PDGFRA+ oligodendrocyte lineage cells in the dorsal pallium of wildtype forebrain slices cultured in DMSO control, 100 ng/mL SHH, and SHH + DAPT, and NICD forebrain slices cultured in 100 ng/mL SHH. Dotted lines indicate the ventral (bottom) limits of the pallium. Scale bar, 100 μm. (D-G) Quantification of oligodendrocyte lineage cells from cultured slices. Total numbers of OLIG2+ cells (D), OLIG2+ PDGFRA-cells (E), and OLIG2+ PDGFRA+ cells (F) were counted per mm^2^ in the pallium of DMSO control, SHH-treated, SHH + DAPT-treated, and NICD + SHH-treated forebrain slice cultures. Graphs show the average ± SEM among biological replicates. Data from (E) and (F) were re-graphed in (G) to show the proportion of total OLIG2+ cells that are either PDGFRA+ or PDGFRA-. One-way ANOVA (with Tukey’s *post hoc*) comparing DMSO, SHH, NICD + SHH: (D) *p*= 2.01e-08; (E and G) *p* = 0.15, n.s. Welch ANOVA (with Games-Howell *post hoc*) comparing DMSO, SHH, SHH + DAPT: (D) *p*= 1.68e-06. One-way ANOVA comparing DMSO, SHH, SHH + DAPT: (E and G) *p*= 0.11, n.s. Kruskal-Wallis (with Dunn’s) comparing DMSO, SHH, NICD + SHH: (F and G) p= 3.07e-05. Kruskal-Wallis (with Dunn’s) comparing DMSO, SHH, SHH + DAPT: (F and G) *p*= 5.23e-05. \**p*< 0.05, \*\*\**p*< 0.0001, ****<0.00001, n.s. = not significant. DMSO control N= 11, SHH N=11, SHH + DAPT N= 6, SHH+NICD N=7. See also Figure S1.

Most OLIG2+ cells in the pallium at this stage are pre-OPCs that are newly derived from dorsal progenitors and do not express PDGFRA yet. Conversely, we previously showed that the majority of OLIG2+ PDGFRA+ cells in the E15.5 pallium are OPCs that derive from the ventral forebrain^5^ and that their numbers are unchanged following DAPT treatment^10^. Similarly, we found here that SHH, SHH+DAPT, and SHH+NICD treatments did not significantly affect OLIG2+ PDGFRA+ OPC numbers in E15.5 slice cultures (**Figures 1C, 1F, and 1G**). On the other hand, SHH treatment increased OLIG2+ PDGFRA-by nearly twofold, and SHH+DAPT treatment again blocked this effect (**Figures 1C and 1E**). We compared the proportion of OLIG2+ cells that are PDGFRA+ versus PDGFRA- and observed that the OLIG2+ PDGFRA-population is the most strongly affected by SHH and DAPT treatments (**Figure 1G**). These results suggest a role for Notch and SHH pathway crosstalk in regulating the initial specification and generation of oligodendrocyte lineage cells from dorsal progenitors.

Neural progenitor responses to SHH signaling depend on the concentration of SHH ligand and duration of exposure^22, 23^. Previous studies in the zebrafish, chick, and mouse spinal cord demonstrated that progenitor responses to a given SHH concentration are potentiated by increased Notch signaling and are decreased when Notch signaling is blocked^11, 12^. This requirement of Notch signaling for optimal SHH-dependent cellular responses can be overcome by increasing SHH signaling directly through its downstream target, Smoothened (SMO)^11^. We therefore wondered whether increasing SHH signaling through SMO activation could promote oligodendrogenesis during Notch inhibition in the dorsal forebrain. We tested this *in vivo* using *in utero* electroporation of dorsal forebrain progenitors (**Figure S1A**). To block Notch signaling we electroporated dominant negative (DN) RBPJ (DN-RBPJ-IRES-mtagBFP2), which inhibits Notch signaling through disruption of the DNA-binding domain of RBPJ^24^. We also generated a plasmid that expresses a constitutively active version of SMO (SMOA1-IRES-GFP), which over-activates the SHH pathway cell-autonomously^25, 26^. We co-electroporated E15.5 mouse embryonic brains with SMOA1 combined with DN-RBPJ or with BFP-only control and analyzed brains at E18.5 by immunohistochemistry for oligodendrocyte lineage markers (**Figure S1A**). As in our prior studies^10^, DN-RBPJ decreased the proportion of electroporated cells that were OLIG2+ and OLIG2+ PDGFRA+ (**Figure S1B-1D**). Conversely, SMOA1 increased the proportion of OLIG2+ cells by almost fourfold (**Figure S1B and S1C**) and OLIG2+ PDGFRA+ OPCs by threefold (**Figure S1B and S1D**). Co-electroporation with SMOA1 and DN-RBPJ together increased OLIG2+ and OLIG2+ PDGFRA+ electroporated cells at similar proportions as SMOA1 alone (**Figure S1B-S1D**). These results indicate that the requirement of Notch signaling for progenitors to produce an optimal SHH response can be bypassed by increasing SHH signaling. Altogether, our data and previous studies demonstrate that Notch signaling can potentiate cellular responses to SHH signaling.

### Notch activation in progenitors promote an undifferentiated transcriptomic identity

We next wanted to understand the molecular mechanisms underlying the ability of Notch signaling to enhance neural progenitor responses to SHH signaling. Following Notch pathway activation, the NICD-RBPJ complex can bind to and activate transcription of many target genes, including Notch transcriptional effectors such as *Hes1*. These changes in gene expression may contribute to the ability of progenitors to generate oligodendrocyte lineage cells in response to SHH. To understand Notch’s influence on gene expression in neural progenitors, we performed bulk mRNA sequencing (RNA-seq) on dorsal forebrain progenitors experiencing Notch overactivation, using the NICD mice. Following dorsal forebrain dissections and dissociations at E16.5, we isolated neural progenitors by magnetic cell sorting (MACS), using an antibody against the progenitor-specific cell-surface marker PROM1^27^ (**Figure 2A**). Three biological replicates were obtained and sequenced for NICD and *Emx1-Cre* only controls (CTRL). Principal component analysis (PCA) of all samples showed separation between CTRL and NICD samples (**Figure 2B**), indicating transcriptomes that were different from one another. Differential gene expression analysis (**Table S1**) revealed 856 upregulated genes in progenitors, including known Notch targets like *Hes1, Hey1*, and *Notch1*. In contrast, 988 genes were downregulated by NICD, including Notch ligands that are known to be expressed in cells with low or no Notch signaling, such as *Dll1* and *Dll3* (**Figure 2C**). Further analysis of differentially expressed genes between NICD and CTRL revealed a transcriptomic profile that was undifferentiated and progenitor-like. For example, genes related to neurogenesis like *Eomes* and *Neurod4* were downregulated in NICD progenitors, whereas stemness genes like *Sox2* and *Nes* were upregulated (**Figure 2C and 2D**). This result is in line with prior studies demonstrating aberrant neurogenesis and increased progenitor maintenance in the *Emx1-Cre;R26-Lox-Stop-Lox-NICD* dorsal forebrain^21^. Interestingly, genes known to play roles in glial differentiation like *Olig2, Pdgfra*, and *Aldh1l1* were also downregulated at this time (**Figure 2D**). Transcriptional targets downstream of the SHH pathway were largely unchanged by NICD overexpression (**Table S2**).

**Figure 2.**
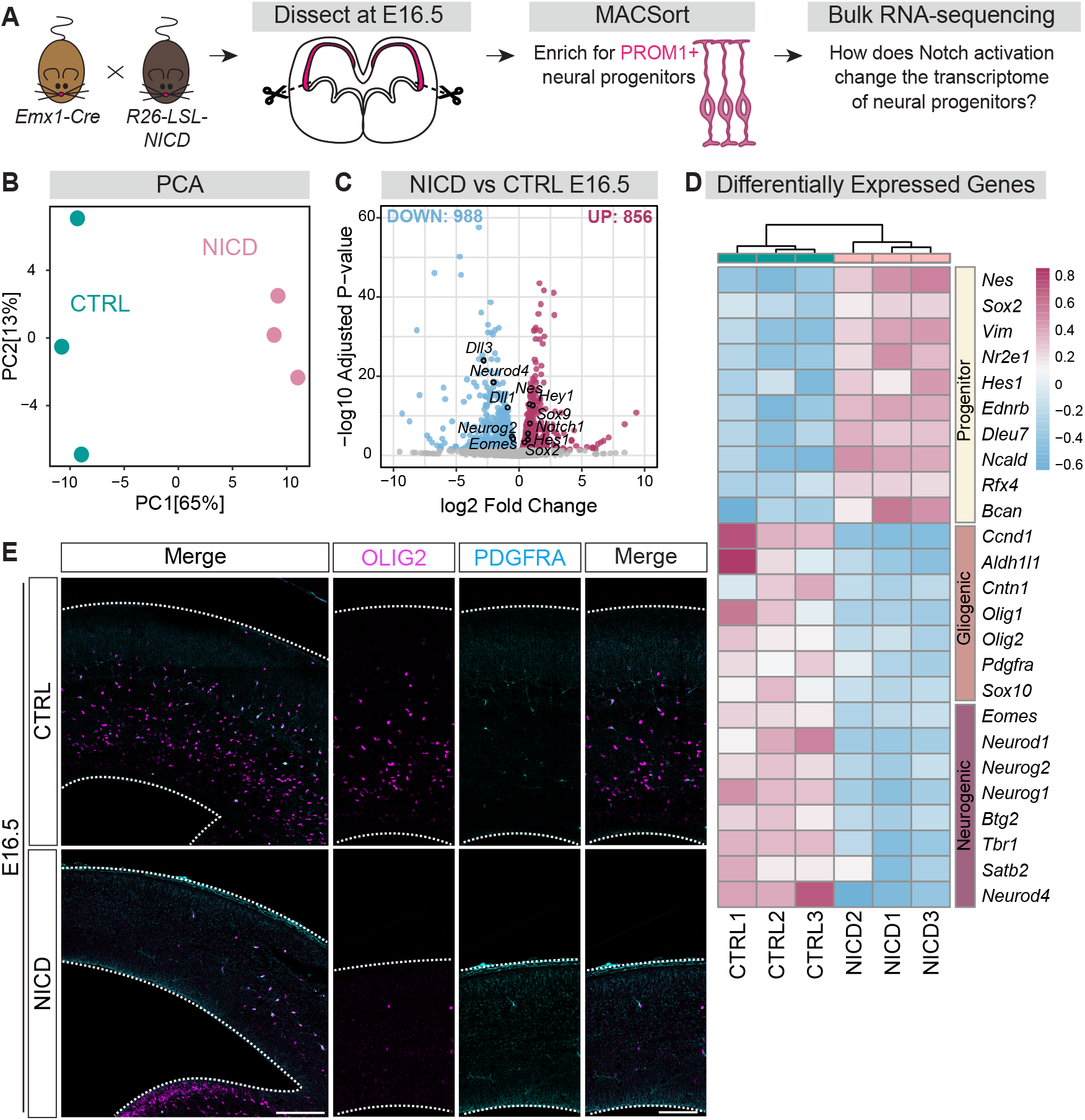
RNA-seq analysis of NICD dorsal progenitors reveals an undifferentiated, stem-like transcriptome. (A) Schematic of bulk RNA-seq workflow. *Emx1-Cre* mice were crossed to *R26-LSL-NICD* mice to activate Notch signaling in *Emx1*+ dorsal progenitors and their progeny. The dorsal forebrains of CTRL and NICD mouse embryos were dissected at E16.5, dissociated, and MACSorted to enrich for PROM1+ dorsal progenitors for bulk RNA-seq. (B) PCA plot indicating separation of CTRL (N= 3) and NICD (N= 3) transcriptomes by condition. PCA = Principal component analysis. (C) Volcano plot representing differentially expressed genes between CTRL and NICD. Differentially expressed genes with adjusted *p* value < 0.05 were considered to be significant. 988 genes were significantly downregulated (blue), and 856 genes were upregulated (pink) in NICD compared to CTRL. The y-axis represents the −log10 Adjusted p-value and the x-axis represents the log2 Fold Change between CTRL and NICD. (D) Heatmap representing selected differentially expressed genes related to progenitor identity (top), gliogenic identity (middle), and neurogenic identity (bottom). Heatmap scale shows variance-stabilized gene expression levels, with colors representing relative expression levels across samples. (E) Representative images of CTRL and NICD E16.5 forebrains, outlined in white, and stained for OLIG2 and PDGFRA. Overview images on the left; scale bar, 200 μm. Zoomed-in images on the right. Dotted lines indicate the dorsal (top) and ventral (bottom) limits of the pallium. Scale bar, 100 μm. See also Tables S1-S2.

Consistent with the decreased expression of glial differentiation genes in the RNA-seq data, we observed very few OLIG2+ or PDGFRA+ cells in the pallium of E16.5 NICD mouse embryos (**Figure 2E**), suggesting that NICD progenitors had not yet begun producing oligodendrocyte lineage cells. This contrasts with CTRL brains that had already initiated oligodendrogenesis at E16.5 (**Figure 2E**). Overall, RNA-seq analysis and IHC at E16.5 demonstrate that NICD-overexpressing progenitors maintained an undifferentiated identity and did not produce OPCs at this time.

### Notch activation changes chromatin accessibility in progenitors

Since NICD dorsal progenitors appeared more progenitor-like and undifferentiated based on transcriptome analyses, we wondered whether Notch’s influence on progenitor responsiveness to SHH and ability to generate the oligodendrocyte lineage may instead be at the level of chromatin accessibility. The chromatin landscape plays key roles in regulating gene expression and thus determining the cell identity of progenitors. To test whether Notch signaling can influence the chromatin landscape in progenitors, we used Assay for Transposase-Accessible Chromatin sequencing (ATAC-seq) to profile accessible chromatin regions in sorted PROM1+ dorsal progenitors (**Figure 3A**). Three biological replicates were obtained for NICD and CTRL dorsal forebrain progenitors at E16.5 and sequenced for accessible chromatin. Similar to our RNA-seq results, NICD and CTRL dorsal progenitors had different chromatin accessibility profiles based on PCA analysis (**Figure 3B**). In particular, open transcription start sites (TSS) were more enriched in NICD than in CTRL (**Figure 3C**). We further analyzed genome-wide differences in chromatin accessibility (**Table S3**) and identified 29,237 sites that increased in accessibility in NICD dorsal progenitors, and 4,731 sites that decreased in accessibility in NICD compared to CTRL progenitors (**Figure 3D**). Since increased chromatin accessibility is associated with a multipotent progenitor state^28^, our ATAC-seq results are in line with our RNA-seq results suggesting that NICD promotes an undifferentiated progenitor identity.

**Figure 3.**
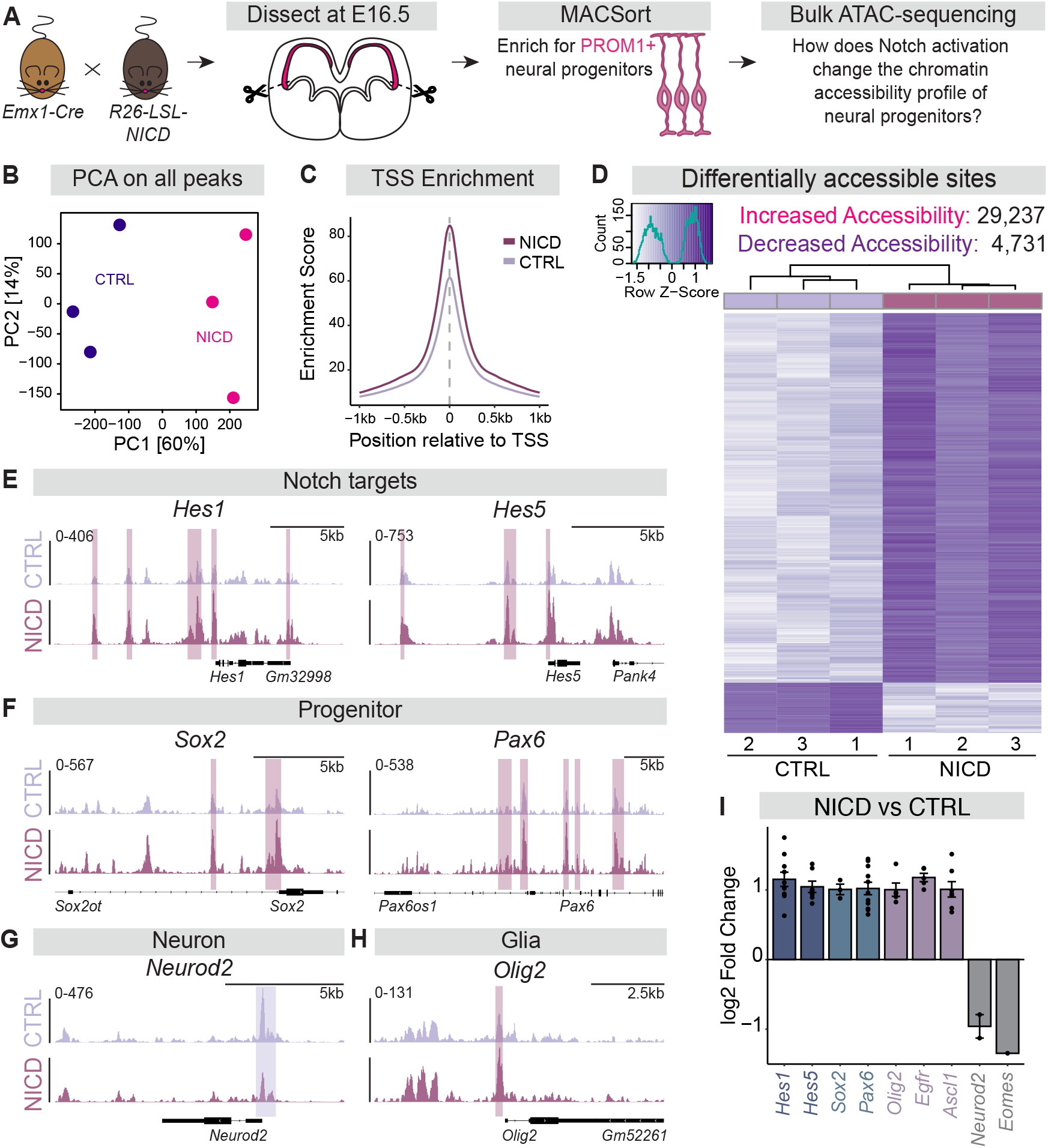
NICD promotes chromatin accessibility in dorsal progenitors as revealed by ATAC-seq. (A) Schematic of bulk ATAC-seq workflow. *Emx1-Cre mice* were crossed to *R26-LSL-NICD* mice to activate Notch signaling in *Emx1*+ dorsal progenitors and their progeny. The dorsal forebrains of CTRL and NICD mouse embryos were dissected at E16.5, dissociated, and MACSorted to enrich for PROM1+ dorsal progenitors for bulk ATAC-seq. (B) PCA plot of all accessible chromatin peaks identified in CTRL (N= 3) and NICD (N= 3) samples. PCA = Principal component analysis. (C) TSS enrichment plot representing normalized read coverage or accessible chromatin around transcription start sites in CTRL and NICD. The y-axis represents an enrichment score or ratio of read density at the TSS compared to background read density. The x-axis represents the position in kb relative to the TSS. TSS= transcription start site; kb = kilobases. (D) Heatmap representing the Z-scores of normalized read counts for all differentially accessible sites that either increased or decreased in NICD compared to CTRL. Peaks are considered significant with an adjusted *p* value < 0.05. The x-axis shown in the scale reflects the row Z-scores, which are based on the standardized deviation of accessibility from the mean, while the y-axis represents the count of peaks. (E-H) Genome tracks of peaks surrounding genes related to the Notch pathway (E), progenitor identity (F), neuron identity (G), and glia identity (H). Peaks highlighted in pink are significant peaks assigned to shown genes and enriched in NICD, while peaks highlighted in purple are significant peaks enriched in CTRL. Scales shown on the y-axis represent signal density across the genome. (I) Graph representing the log2 Fold Change of all significant peaks associated with the list of selected genes, with each dot representing a peak. Peaks were assigned to genes based on distance to the nearest TSS. See also Table S3.

To further understand the influence of NICD on cell identity, we analyzed genes associated with differentially accessible regions. Notch target genes like *Hes1* and *Hes5* exhibited chromatin peaks that were significantly more accessible in NICD compared to CTRL (**Figures 3E and 3I**). Similarly, progenitor identity genes, like *Sox2* and *Pax6*, also showed increased accessibility in NICD progenitors (**Figures 3F and 3I**). NICD progenitors had decreased chromatin accessibility peaks near certain neurogenic genes like *Neurod2* and *Eomes* (**Figures 3G and 3I**), consistent with the RNA-seq results indicating a less neurogenic state. Interestingly, however, peaks near genes associated with gliogenesis like *Olig2, Egfr*, and *Ascl1* (**Figures 3H and 3I**) exhibited increased accessibility. These results indicate that Notch activation with NICD reorganizes chromatin near lineage specification genes. We therefore wanted to better understand the overall influence of NICD on the chromatin landscape surrounding broader categories of genes associated with different cell fates. To do so, we curated gene lists known to be important for progenitor, neurogenic, and gliogenic identities, and compared the normalized cumulative accessibility of chromatin near the genes in each identity class. Interestingly, NICD-overexpressing progenitors exhibited abundant accessibility near all three gene categories (**Figure 4A-4C**), as well as genes specific to the oligodendrocyte lineage (**Figure 4D**). Therefore, whereas the transcriptomic profile of NICD progenitors indicated an undifferentiated and progenitor-like state, their chromatin accessibility profile suggested that they had increased accessibility nearby both neurogenic and gliogenic genes that were not yet expressed. Together with our RNA-seq data, our ATAC-seq analyses point towards the possibility that these genes with increased accessibility are poised to be expressed but require specific factors that are not yet present. Such factors may be transcriptional regulators that play key roles in determining cellular identity.

**Figure 4.**
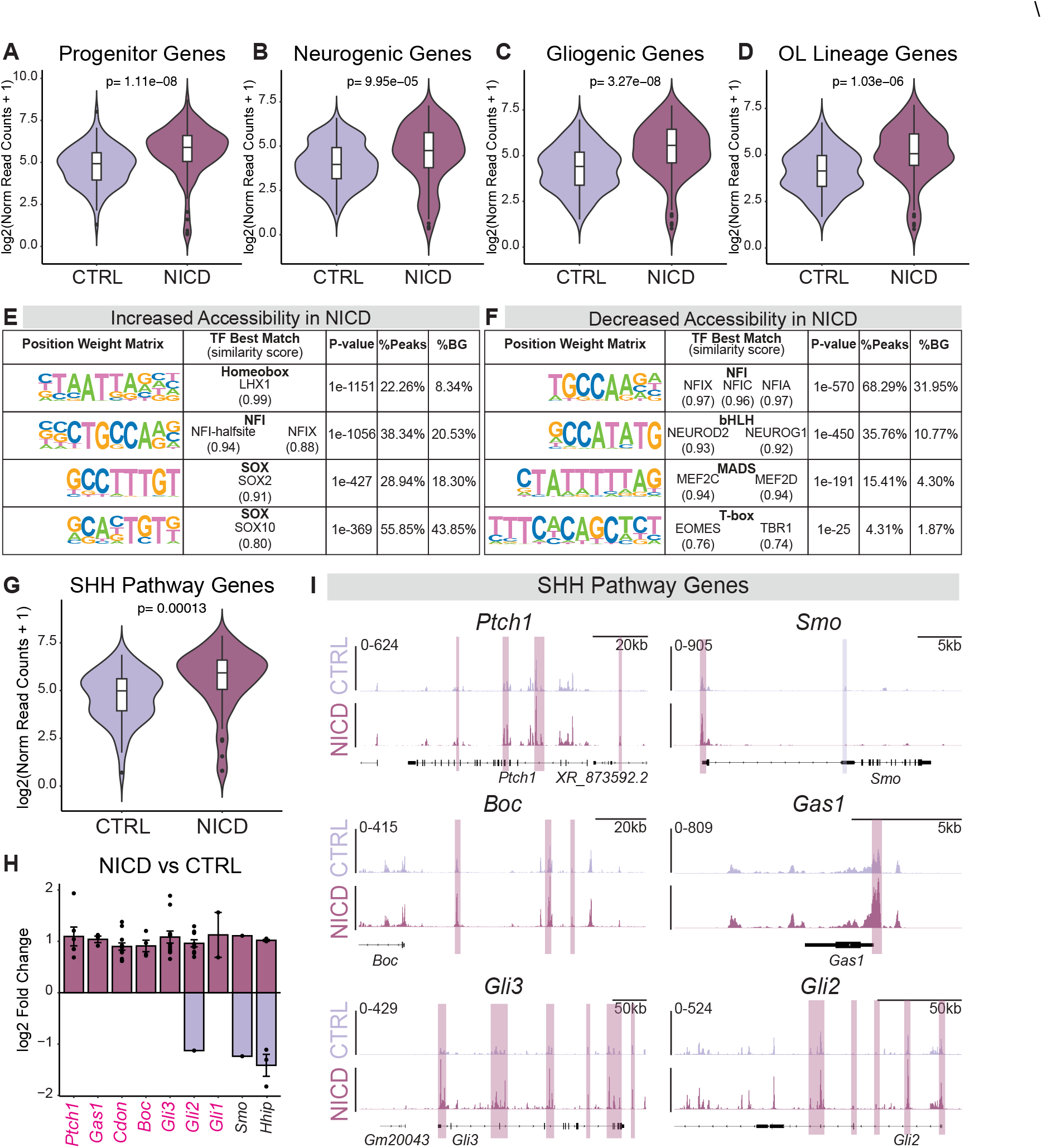
Notch over-activation reorganizes chromatin surrounding cell identity genes and SHH pathway genes. (A-D) Plots representing the normalized read counts or accessibility of significant peaks assigned to progenitor genes (A), neurogenic genes (B), gliogenic genes (C), OL lineage genes (D). Wilcoxon rank-sum test was used to determine significance. OL = oligodendrocyte lineage. (E-F) Transcription factor binding motifs identified by HOMER *de novo* motif enrichment analysis of the 29,237 increased accessibility sites (E) and 4,731 decreased accessibility sites (F) in NICD. % BG = % of Background sequences. (G) Normalized read count plot of significant peaks assigned to genes related to the SHH signaling pathway. (H) Graph representing the log2 Fold Change of all significant peaks associated with selected SHH pathway genes, with each dot representing a peak. Peaks were assigned to genes based on distance to the nearest TSS. TSS = Transcription Start Site. Gene names colored in pink have peaks that contain RBPJ binding motifs. (I) Genome tracks showing chromatin accessibility peaks surrounding genes related to the SHH pathway. Peaks highlighted in pink are significant peaks assigned to the shown genes and enriched in NICD, while peaks highlighted in purple are significant peaks enriched in CTRL. See also Tables S4-S5.

Therefore, we next asked which transcription factor motifs were present in both more accessible and less accessible sites in NICD progenitors. *De novo* motif enrichment analysis of more accessible sites predicted motifs of transcription factors like LHX family, NFI family, and SOX10 (**Figure 4E**), all of which have known roles in gliogenesis in the forebrain^29–32^. The motif that matches SOX2, which influences progenitor identity^33, 34^, was also identified (**Figure 4E**). In contrast, the motifs identified in less accessible sites (i.e., enriched in CTRL) matched with transcription factors like NEUROD2, MEF2C, and EOMES (**Figure 4F**), which are known regulators of neuron identity^35–38^. Furthermore, these motifs were identified near genes associated with neuron differentiation and maturation (**Table S4**). Since these motifs were found in sites that are less accessible in NICD-overexpressing progenitors, this suggests that NICD may prevent neuron differentiation and maturation by decreasing chromatin accessibility to neurogenic transcription factors. On the other hand, LHX, NFI, and SOX10 motifs were enriched in regions with increased accessibility in NICD progenitors and were found near genes important for gliogenesis (**Table S5**). Interestingly, motifs matching the binding motifs of the NFI family of transcription factors were identified in both more and less accessible sites, consistent with their known roles in both neurogenesis and gliogenesis^39^. Altogether our ATAC-seq and RNA-seq data from E16.5 dorsal forebrain progenitors indicate that Notch signaling reinforces progenitor stemness at the genomic and transcriptional levels, while also epigenetically priming progenitors to turn on gliogenic genes that are not yet expressed.

### Notch activation promotes chromatin accessibility nearby SHH pathway component genes

Since Notch signaling is required for SHH-induced oligodendrogenesis and NICD promoted chromatin accessibility near oligodendrocyte fate specification genes, we next asked whether Notch activation can regulate the chromatin landscape surrounding genes known to promote SHH signaling. We compared the normalized accessibility of peaks associated with genes that positively regulate SHH pathway activity between CTRL and NICD. We found that NICD had increased accessibility near SHH pathway genes (**Figure 4G**). When we compared the log2 fold change of accessibility at all peaks that were associated with SHH pathway component genes, we found that almost all chromatin accessibility changes were in the positive direction (**Figure 4H-I**). Although the RBPJ motif was not identified in the motif enrichment analysis, a search for instances of the RBPJ motif in our ATAC-seq data revealed that several differential chromatin peaks associated with SHH pathway component genes contained RBPJ motifs, suggesting the possibility that NICD-RBPJ may be regulating some of these genes directly (**Figure 4H**, highlighted in pink). Altogether, these ATAC-seq analyses suggest that NICD promotes chromatin accessibility nearby SHH pathway component genes.

### Notch signaling primes progenitors for enhanced SHH signaling and robust oligodendrogenesis

Our results indicate that Notch signaling may be priming neural progenitors epigenetically for enhanced SHH signaling and gliogenesis, while also transcriptionally repressing glial and neuronal differentiation. This is consistent with our prior studies demonstrating that Notch signaling plays dual roles in setting the timing of the neuron-glia switch by both positively and negatively regulating oligodendrogenesis^10^. We therefore hypothesized that NICD-overexpressing progenitors that are epigenetically primed but transcriptionally repressed at E16.5 would exhibit enhanced SHH signaling and robust oligodendrogenesis at E17.5, which is the peak of the SHH-dependent neuron-glia transition ^5^. To test this hypothesis, we first stained sections from NICD and CTRL brains with oligodendrocyte lineage markers at E17.5 (**Figure 5A**). In contrast to the lack of OLIG2+ and OLIG2+ PDGFRA+ cells in NICD brains at E16.5 (**Figure 2E**), we observed robust oligodendrogenesis in NICD-overexpressing brains at E17.5 (**Figure 5B-C**). We quantified the density of OLIG2+ and OLIG2+ PDGFRA+ cells in the ventricular and subventricular zones (VZ/SVZ), where they are first specified and generated, and found a significant increase in NICD-overexpressing brains compared to CTRL brains (**Figures 5D and 5E**). These data suggest that increased Notch signaling in NICD brains delayed oligodendrogenesis at E16.5, but enhanced oligodendrocyte lineage output one day later at E17.5.

**Figure 5.**
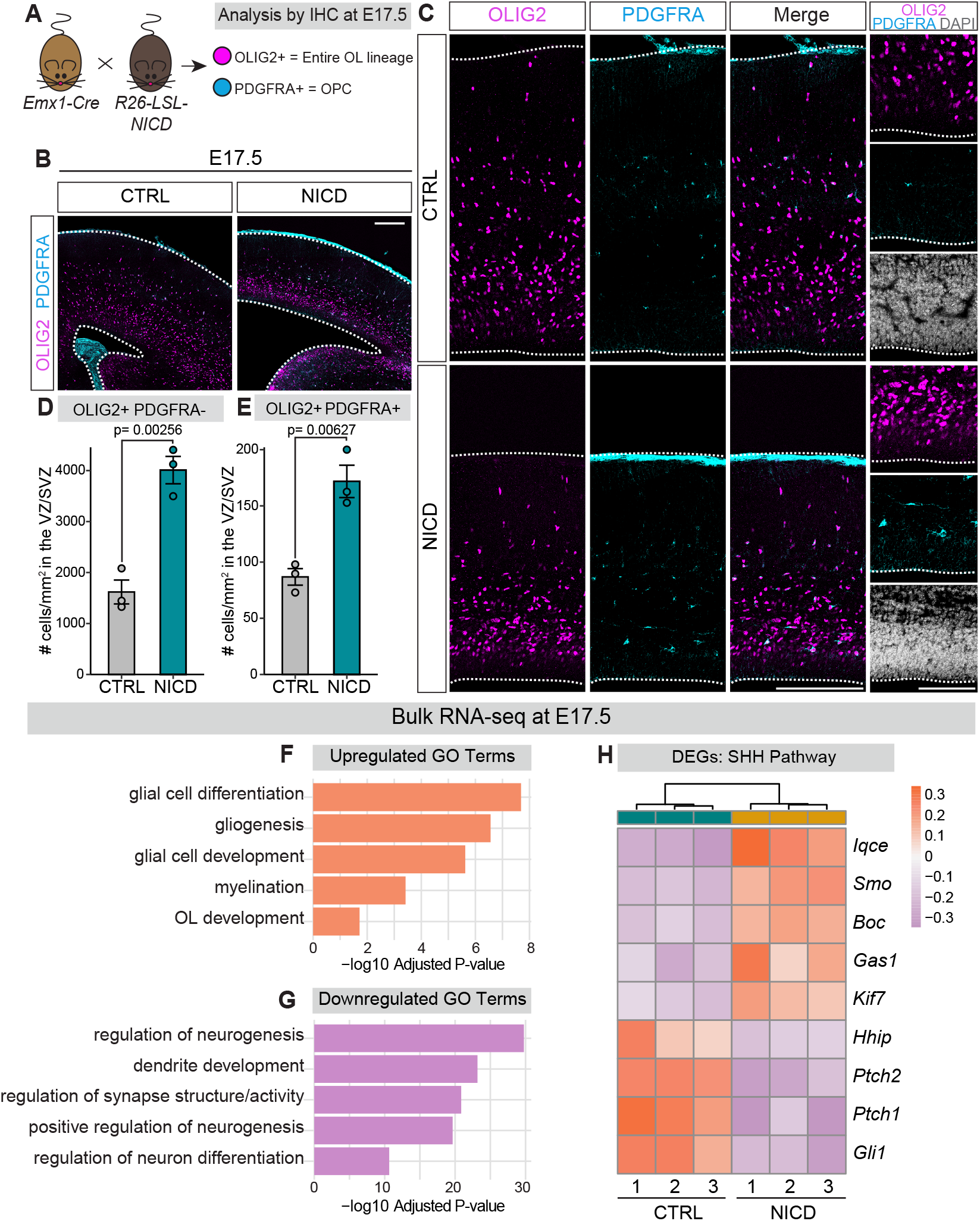
NICD brains have increased gliogenesis and upregulated expression of SHH pathway components at E17.5. (A) Schematic of workflow and IHC. *Emx1-Cre* mice were crossed to *R26-LSL-NICD* mice to activate Notch signaling in *Emx1*+ dorsal progenitors and their progeny. Brains were dissected at E17.5 and stained for OL markers. OL = oligodendrocyte; OPC = oligodendrocyte precursor cell. IHC = immunohistochemistry. (B) Representative overview images of CTRL and NICD E17.5 forebrains, outlined in white, and stained for OLIG2 and PDGFRA. Scale bar, 200 μm. (C) On the left, representative images of CTRL and NICD dorsal palliums. Dotted lines indicate dorsal (top) and ventral (bottom) limits. Scale bar, 100 μm. Zoomed-in images on the far right show the VZ/SVZ where dorsal progenitors reside. Scale bar, 100 μm. (D-E) Quantification of # of OLIG2+ PDGFRA-(D) and OLIG2+ PDGFRA+ (E) cells per mm^2^ within the VZ/SVZ +/-SEM. VZ/SVZ = ventricular/subventricular zones. (F-G) GO enrichment analysis on upregulated (F) and downregulated (G) genes identified by RNA-seq. X-axis represents the −log10 Adjusted *P* value, with GO terms on the y-axis. GO = gene ontology. (H) Heatmap representing selected differentially expressed genes related to the SHH pathway. Heatmap scale shows variance-stabilized gene expression levels, with colors representing relative expression levels across samples. DEG = differentially expressed genes. See also Figure S2 and Table S6.

We next repeated our bulk RNA-seq experiments of PROM1+ isolated neural progenitors, this time from E17.5 NICD and CTRL brains. NICD and CTRL samples clearly separated in our PCA analysis, indicating divergent transcriptomes (**Figure S2A**). Differential gene expression analysis (**Table S6**) revealed 3,982 genes that were upregulated and 4,040 genes that were downregulated by NICD at E17.5 (**Figure S2B**). We performed gene-ontology (GO) enrichment analysis separately for both upregulated and downregulated differentially expressed genes. GO terms that were enriched based on the significantly upregulated genes included “glial cell differentiation”, “gliogenesis”, “myelination”, as well as “OL development” (**Figure 5F**). Conversely, downregulated GO terms included “positive regulation of neurogenesis”, “dendrite development”, and “regulation of neuron differentiation”, indicating suppression of neuron differentiation and maturation (**Figure 5G**). In addition, many of the gliogenic transcription factors whose binding motifs were enriched within increased accessibility chromatin regions in ATAC-seq data from E16.5 also exhibited increased expression in RNA-seq data by E17.5, whereas expression of neurogenic transcription factors remained low (**Figure S2D-E**). Together, these results support that increased Notch signaling enhances gliogenesis at E17.5, while repressing neuron differentiation transcriptional programs.

Given that SHH is a key regulator of oligodendrogenesis, we investigated whether Notch signaling influences SHH pathway gene expression. GO enrichment analysis of upregulated genes highlighted “smoothened signaling pathway” as one of the most enriched terms (**Figure S2C**), referring to the signal transduction process involving the SHH pathway component SMO^40^. Further examination of NICD differentially expressed genes revealed upregulation of *Smo*, SHH co-receptors *Boc* and *Gas1*, as well as cilia-associated genes *Iqce* and *Kif7* (**Figure 5H**), all of which are known to positively regulate SHH signaling^40–45^. Additionally, NICD-overexpressing brains showed significant decreases in the expression of known SHH pathway inhibitors, including *Hhip, Ptch2*, and *Ptch1* (**Figure 5H**)^46–50^. *Gli1*, a known downstream target and activator of SHH signaling^51, 52^, was decreased in NICD-overexpressing brains (**Figure 5H**). Furthermore, GO enrichment analysis of upregulated genes revealed terms such as “cilium organization”’ and “cilium assembly” (**Figure S2C**), suggesting that NICD may enhance primary cilium structure, which is critical for the proper activation of SHH signaling^45, 53, 54^. Overall, our RNA-seq analysis at E17.5 demonstrated that increased Notch signaling prepares dorsal progenitors to respond to SHH by regulating components involved in SHH signal transduction and ciliary structure. This suggests that Notch signaling primes these progenitors for increased sensitivity and responsiveness to SHH signaling.

Given the influence of Notch signaling on the expression of SHH pathway components, we asked how manipulating Notch signaling up or down in neural progenitors would affect SHH signaling strength at E17.5. We employed a GFP reporter under control of GLI binding sites (GLIBS-GFP) that has been used to measure SHH transcriptional activity^55^. To test whether Notch signaling affects SHH transcriptional activity *in vivo*, we co-electroporated GLIBS-GFP with either DN-RBPJ-IRES-mTagBFP2, NICD-IRES-mTagBFP2, or BFP only control at E15.5 and analyzed brains for reporter activity at E17.5 (**Figure 6A**). We simultaneously monitored Notch signaling activity by co-electroporating a Notch transcriptional reporter under control of the *Hes5* promoter (*Hes5*-dsRed)^56^. In BFP control brains, some electroporated BFP+ progenitors within the VZ exhibited signal from both the *Hes5*-dsRed and GLIBS-GFP reporters (green arrows), whereas others lacked signal from either reporter (blue arrow) (**Figure 6B**). Measuring the fluorescence intensity of the GLIBS-GFP and *Hes5*-dsRed reporters revealed that they were positively and linearly correlated in individual BFP+ progenitors, such that progenitors with higher *Hes5*-dsRed levels also had higher GLIBS-GFP levels (**Figure 6C**). As expected, DN-RBPJ significantly reduced the fluorescence levels of *Hes5*-dsRed, confirming DN-RBPJ’s inhibitory effects on Notch signaling activity (**Figure 6B and 6D**). We also observed that GLIBS-GFP signal was almost completely gone in DN-RBPJ-electroporated brains (**Figure 6B and 6E**), indicating that Notch pathway inhibition blocked SHH transcriptional activity in dorsal progenitors. Conversely, NICD overexpression increased the fluorescence intensity of both *Hes5*-dsRed and GLIBS-GFP (**Figure 6B, 6D, and 6E)**. Therefore, Notch activation is sufficient to increase the levels of SHH transcriptional activity in dorsal progenitors. Altogether, these results indicate that Notch signaling regulates the response of progenitors to endogenous SHH signaling in the dorsal forebrain.

**Figure 6.**
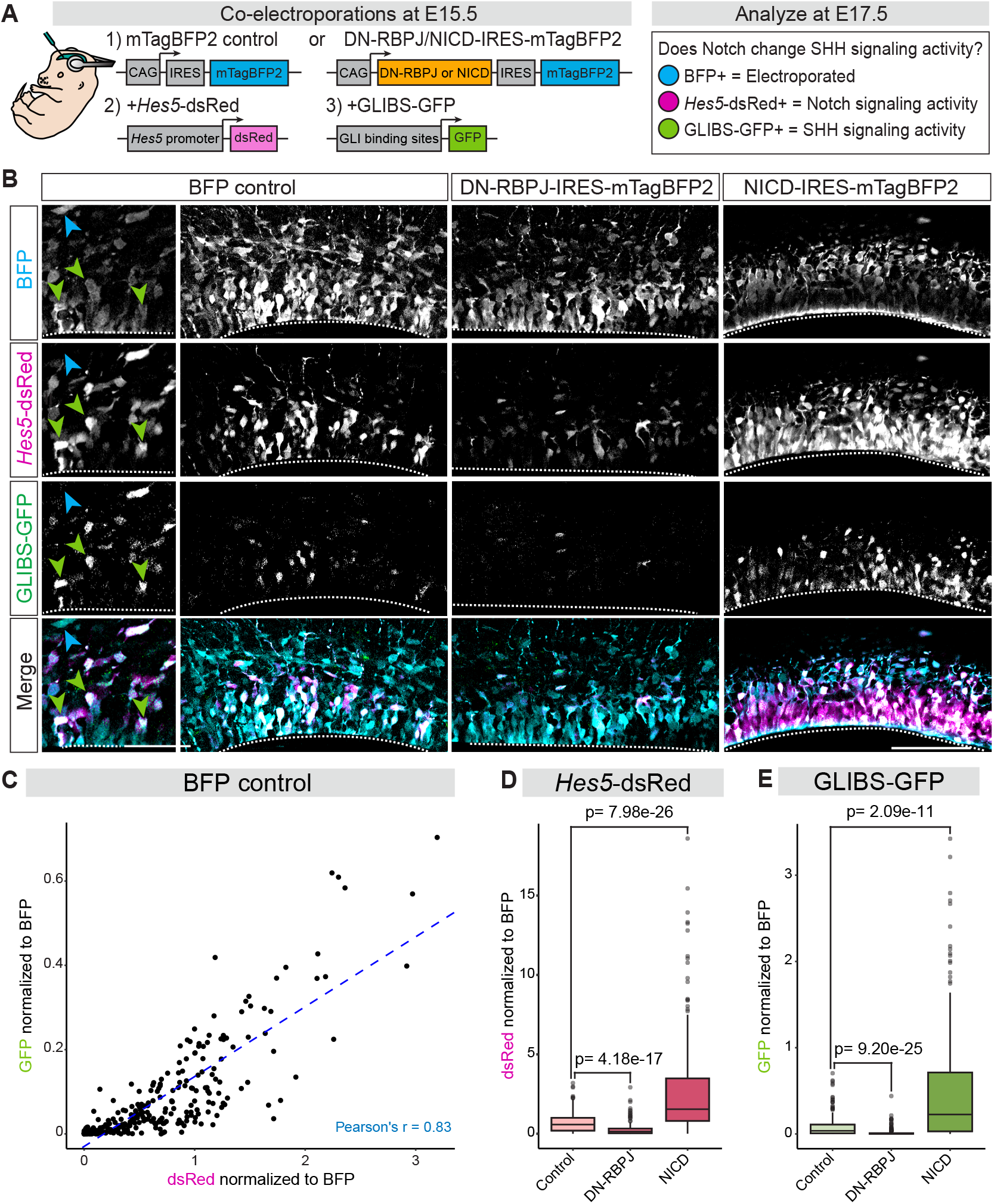
Notch signaling regulates SHH activity reporter levels in dorsal progenitors in vivo. (A) Schematic of constructs for BFP control, DN-RBPJ-IRES-mTagBFP2, NICD-IRES-mTagBFP2, *Hes5*-dsRed, and GLIBS-GFP. Wildtype mouse embryos were co-electroporated at E15.5 and analyzed for Notch and SHH reporter activity at E17.5. (B) Representative images of brains electroporated with BFP control, DN-RBPJ-IRES-mTagBFP2, or NICD-IRES-mTagBFP2 along with Notch (*Hes5*-dsRed) and SHH (GLIBS-GFP) activity reporters. Left panels in BFP control are zoomed in images showing BFP+ cells that are *Hes5*-dsRed and GLIBS-GFP double positive (green arrows) and a BFP+ cell that is double negative for both reporters (blue arrow). Scale bar, 50 μm. Representative overview images of *Hes5*-dsRed and GLIBS-GFP fluorescence in the ventricular zone of BFP control, DN-RBPJ, and NICD brains. Dotted lines outline the ventral (bottom) limits of the pallium. Scale bar, 100 μm. Images for the GFP channel were adjusted with increased brightness for the purpose of visibility in the figure. (C) Graph of the normalized fluorescence intensity of dsRed plotted against the normalized fluorescence intensity of GFP, with each dot representing an individual cell in BFP control brains. The blue dotted line represents the line of best fit. The Pearson’s *r* value represents the strength of linear correlation. (D-E) Quantification of the normalized fluorescence intensity of *Hes5*-dsRed (D) or GLIBS-GFP (E) in Control, DN-RBPJ, and NICD (± SEM among biological replicates). Kruskal-Wallis test *p*<2.2e-16 for (D) and (E). Dunn’s test to test significance between conditions, with p values adjusted using the Bonferroni method. BFP control N= 267 cells, DN-RBPJ N= 285 cells, NICD N= 290 cells.

## DISCUSSION

During embryogenesis, multipotent progenitor cells are directed toward distinct cell fates at specific developmental stages through a combination of intrinsic genetic programs and external signaling cues^2, 9, 57, 58^. How do the intrinsic properties of progenitor cells shape their interpretation of external signals and the specific cell identities they adopt in response to those signals? To address this, we focused on the neuron-glia switch in the developing dorsal forebrain; a process in which neural progenitors transition from producing neurons to oligodendrocytes in response to a late embryonic SHH signal ^5^. We previously identified Notch signaling as an important regulator of the timing of this switch^10^, but the downstream mechanisms leading to the acquisition of oligodendrocyte fates remained unclear.

Here, we now show that Notch signaling regulates the progenitor response to SHH, thereby facilitating the neuron-glia transition. We propose that Notch signaling establishes an internal progenitor state through three key mechanisms: 1) increased Notch activity transcriptionally prolongs progenitor maintenance and prevents differentiation, 2) Notch signaling epigenetically and transcriptionally primes progenitors to respond efficiently to SHH, and 3) Notch signaling establishes a chromatin landscape that suppresses neuronal fates while priming progenitors for gliogenesis. These findings support a working model in which Notch signaling establishes progenitor sensitivity to SHH and competence for oligodendrocyte fates at both the epigenetic and transcriptional levels, permitting the timely production of oligodendrocytes when SHH signaling peaks in the dorsal forebrain (**Figure 7**).

**Figure 7.**
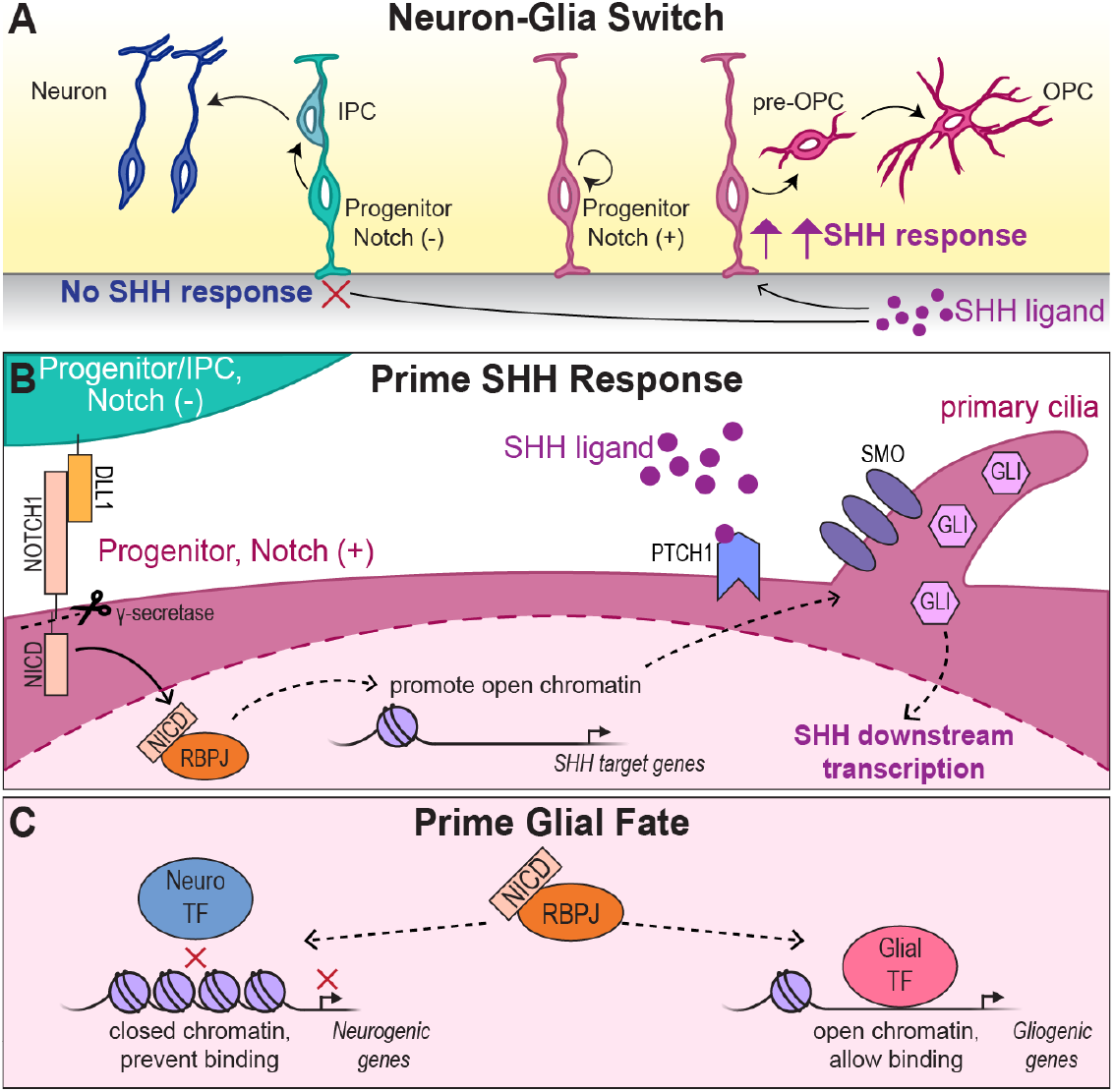
Proposed model for Notch signaling in regulating SHH response and glial fate specification. (A) During the neuron-glia switcn, dorsal forebrain progenitors with low or no Notch activity do not respond to SHH ligand and differentiate into intermediate progenitor cells (IPC), which generate upper-layer excitatory neurons. Meanwhile, higher Notch signaling in other progenitors inhibits neurogenesis and maintains the progenitor state. These Notch (+) progenitors have increased sensitivity to SHH ligand, which initiates production of pre-OPCs and OPCs. (B) Notch activity primes dorsal forebrain progenitors to respond to SHH. The Notch ligand, DLL1, from Notch (−) cells binds to the NOTCH1 receptor of neighbonng Notch (+) cells, leading to the cleavage of NICD by ⊠-secretase. NlCD translocates to the nucleus and forms a transcriptional complex with RBPJ to activate transcription of unknown factors that open chromatin near SHH target genes, thereby enhancing SHH signaling and downstream transcription. (C) NICD-RBPJ complex reduces chromatin accessibility near neurogenic genes and prevents binding to neurogenic transcription factors. NICD-RBPJ also promotes chromatin opening near gliogenic genes, allowing transcription factors to bind and promote glial fates.

### Notch signaling regulates the response of dorsal forebrain progenitors to SHH

SHH signaling activity is initially low in the dorsal forebrain during early embryogenesis when neurogenesis is actively occurring. However, during later developmental stages, SHH signaling increases and triggers the shift to oligodendrocyte fate specification^5^. This rise in SHH signaling is facilitated by ligand secretion from migrating interneurons and the choroid plexus^5^. In our current study, *ex vivo* slice culture experiments revealed that Notch inhibition effectively blocked SHH-induced production of OLIG2+ cells, suggesting that Notch signaling is necessary for progenitors to respond to SHH and produce glia. Several studies have demonstrated significant crosstalk between the Notch and SHH pathways in various contexts, including the developing spinal cord and retina^11–16^. Notably, in the developing mouse and chick spinal cord, Notch activity regulates progenitor sensitivity to SHH. Consistent with this, we found that Notch over-activation in dorsal forebrain progenitors enhanced SHH signaling activity, whereas Notch inhibition nearly abolished it. Given that SHH ligand and downstream signaling components are expressed at lower levels in the dorsal forebrain compared to more ventral regions of the central nervous system^59–61^, Notch-dependent priming of progenitors for SHH responsiveness may be particularly important during the neuron-glia switch in the dorsal forebrain (**Figure 7A**). However, cooperation of Notch and SHH during gliogenesis may be a more broadly conserved mechanism throughout the CNS and across species. Notch signaling is required for oligodendrocyte fate specification in the zebrafish spinal cord, where it maintains subsets of neural progenitors during neurogenesis and specifies them toward glial fates^62^. Interestingly, constitutive Notch activity promotes excess OPCs in the zebrafish spinal cord, but only at the place and time when OPCs normally form, suggesting that other signals like SHH are also required^62^. In line with this idea, the Notch and SHH pathways have been shown to cooperate to drive oligodendrogenesis in the zebrafish spinal cord^12^.

Signaling crosstalk between Notch and SHH occurs at multiple levels, from transcriptional control to protein activity, and varies with biological context^11, 13–16, 63^. In spinal cord progenitors, Notch signaling modulates the localization of key SHH pathway components within primary cilia, a critical site for SHH signal transduction. Notch activation prevents PTCH1 from localizing to the primary cilia, thereby allowing the accumulation of SMO and subsequent activation of GLI transcription factors^11, 13^. Here, we provide evidence for an epigenetic regulatory mechanism in which Notch signaling modulates chromatin accessibility at SHH pathway genes in dorsal forebrain progenitors (**Figure 7B**). Increased Notch signaling facilitated chromatin accessibility associated with genes encoding SHH pathway components. The presence of RBPJ binding motifs at chromatin peaks associated with SHH pathway genes suggests a potential direct regulatory role for Notch, consistent with a role for RBPJ in chromatin modulation through regulation of histone modifications in other contexts^64^. Remarkably, expression of most SHH pathway components was initially unchanged at E16.5 but were more highly expressed in NICD-overexpressing progenitors one day later. Increased accessibility and higher expression of *Smo, Boc*, and *Gas1* and reduced expression of *Ptch1* and *Hhip* suggests that Notch activation may promote SHH pathway responsiveness through the upregulation of co-receptors while sustaining pathway activity by repressing inhibitors. Additionally, our reporter assays demonstrated increased GLI-dependent transcriptional output even though *Gli2* and *Gli3* mRNA expression levels remained unchanged, further suggesting that Notch signaling impacts SHH signaling upstream of GLI transcription factors. Interestingly, GO analysis highlighted significant enrichment for terms related to cilium structure and organization. *Kif7*, a regulator of GLI proteolytic processing in the cilium^43, 44^, and ciliary function genes such as *Iqce, Evc*, and *Evc2*^45, 65^ were significantly upregulated. Given the importance of primary cilia in SHH signal transduction and that Notch signaling can promote longer cilia^13, 53, 54^, it is plausible that Notch signaling contributes to ciliary integrity as part of the mechanism for activating and sustaining SHH signaling (**Figure 7B**).

### Notch-mediated epigenetic priming guides progenitor competence and cell fate decisions

Epigenetic priming is a process by which cells modify chromatin structure to make specific genes more accessible to regulatory factors, thereby facilitating gene expression and guiding cell fate transitions^66, 67^. Our findings reveal that Notch signaling plays an important role in epigenetic priming of dorsal forebrain progenitors toward glial fates. ATAC-seq analysis of NICD-overexpressing progenitors showed increased chromatin accessibility at genes associated with progenitor, gliogenic, and neurogenic identities, suggesting a state of multilineage potential. In support of this, multilineage priming has been described in early embryonic dorsal progenitors, where expression of diverse neuronal subtype markers reflects a mixed transcriptional identity before these progenitors commit to specific fates^8^. Our RNA-seq analysis of NICD-overexpressing progenitors at E16.5 showed that progenitor-associated genes were highly expressed, whereas differentiation-specific mRNAs remained low. This result indicates an undifferentiated state in which these genes are epigenetically poised, but not yet expressed.

Despite this multilineage potential, further ATAC-seq analysis revealed that the resulting chromatin landscape favored gliogenesis through two main mechanisms. First, increased Notch signaling promoted chromatin accessibility at gliogenic genes, including those associated with the oligodendrocyte lineage (**Figure 7C**). Second, Notch activation reduced accessibility at regions containing neurogenic transcription factor binding sites, while increasing accessibility at regions enriched with binding sites for gliogenic transcription factors (**Figure 7C**). Notch signaling is well-documented for its role in inhibiting neurogenesis by repressing the transcription of pro-neural factors^68–70^, a pattern we observed in our RNA-seq data. However, our findings suggest an additional mechanism by which Notch suppresses neurogenesis. By reducing chromatin accessibility at neurogenic regulatory sites, Notch activation may prevent transcription factor binding and activation of neurogenic programs. Conversely, increased accessibility for gliogenic transcription factor motifs suggests that epigenetic priming by Notch signaling predisposes progenitors toward glial fates (**Figure 7C**). In fact, the gliogenic transcription factors whose binding motifs were enriched within more accessible chromatin regions also exhibited increased expression from E16.5 to E17.5, whereas neurogenic transcription factor expression remained low. This idea of Notch priming glial fates is further supported by our findings that by E17.5, NICD-overexpressing brains exhibited an enrichment of glial-related GO terms in RNA-seq data as well as increased production of OPCs. In the future, it will be interesting to determine what NICD-RBPJ targets directly modify chromatin to allow changes in accessibility and subsequently transcription.

### Changes in Progenitor Competence over Developmental Time

Previous studies have shown that progenitors undergo developmental changes over time, progressively maturing while maintaining a core progenitor identity defined by stable expression of progenitor markers. However, as they mature, their expression of other genes changes, which may alter progenitor competence to developmental cues at different times^6, 71–73^. This progressive change in competence suggests that progenitors might also feature a flexible chromatin landscape that facilitates these changes while preserving their progenitor state. Supporting this idea, dorsal progenitors have been shown to dynamically remodel their chromatin structure to determine whether they generate neurons or astrocytes in response to BMP signaling^7^. Given the Notch pathway’s established role in maintaining the progenitor pool throughout brain development^74, 75^, it is conceivable that Notch not only preserves progenitor identity but also facilitates chromatin modifications that enable progenitors to respond specifically to developmental cues as they arise. We propose that Notch signaling primes a chromatin landscape in dorsal progenitors that is both poised for gliogenic fates and responsive to SHH, ultimately promoting the specification and generation of glial lineages. This complex interplay underscores how Notch signaling orchestrates both immediate and future gene regulatory programs, and positions progenitors to transition efficiently toward glial fates in response to SHH while also preserving the progenitor pool.

## METHODS

### Experimental model details

*Mouse lines*. The following mouse lines were obtained from The Jackson Laboratory: B6 (C57BL/6J, stock no. 000664); R26-LSL-NICD (*Gt(ROSA)26Sor*^*tm1(Notch1)Dam*^/J (ROSA26^loxP-stop-loxP-Notch1-ICD^), stock no. 008159); Emx1-Cre (B6.129S2-Emx1tm1(cre)Kri/J, stock no. 005628). We also obtained timed pregnant Crl:CD1(ICR) mice (strain no. 022) from Charles River, which arrived at embryonic day (E) 12.5 and on which we performed electroporations at E15.5. For timed matings of B6 wildtype mice, we considered embryos to be at gestation day 0.5 on the day when the vaginal plug was detected. *Emx1-Cre (cre/cre)* mice were crossed with *R26-LSL-NICD (flox/flox)* mice to generate pregnant dams carrying double heterozygous embryos for ATAC-seq, RNA-seq, and immunohistochemistry experiments at either E16.5 or E17.5. *Emx1-Cre* mice crossed with B6 mice were used as controls. Mouse lines were authenticated via genotyping by PCR. Animals were maintained according to the guidelines from the Institutional Animal Care and Use Committee of the University of Colorado-Anschutz Medical Campus. Sex of embryos was not determined for experiments.

## Method details

### Expression plasmids

A *piggyBac* (PB) transposase system was used for *in utero* electroporation experiments to permit stable integration of the reporter plasmid into electroporated progenitors^88^. We found this approach to be necessary to label the oligodendrocyte lineage, in line with previous reports that episomal plasmids are silenced or lost in glial lineages^89, 90^.

DNA fragments for cDNA encoding a constitutively active version of mouse Smoothened (SMOA1) containing the point mutation W539L was synthesized and purchased from IDT as a gBlock. SMOA1 cDNA gBlock was cloned using the NEBuilder HiFi DNA Assembly Master Mix into a *piggyBac* (pPB) expression vector containing a CMV promoter and CAG enhancer, and IRES-GFP (pPB-CAG-IRES-GFP). Generation of plasmids expressing NOTCH1 Intracellular Domain (NICD: AA 1748-2293) and the dominant-negative version of RBPJ (DN-RBPJ) were described previously (Tran et al., 2023). The NICD and DN-RBPJ cDNA gBlocks were cloned into a pPB-CAG-IRES-mTagBFP2 backbone, in which GFP was replaced by mTagBFP2. pPB-CAG-IRES-GFP and pPB-CAG-IRES-mTagBFP2 empty vectors was used as the GFP and BFP controls, respectively. All constructs were confirmed by restriction digest and whole-plasmid DNA sequencing (plasmidsaurus). The PBase expression plasmid CMV-mPB was described previously^5^.

For reporter assays, a plasmid expressing GFP under control of GLI binding sites (pGL3b-8xGliBS:EGFP; GLIBS-GFP) was used to analyze SHH signaling. pGL3b-8xGliBS:EGFP (Addgene plasmid #84602) was a gift from James Chen^55^. To analyze Notch signaling, we used a plasmid expressing dsRed under control of the *Hes5* promoter (*Hes5p*-DsRed; *Hes5*-dsRed). Hes5p-DsRed (Addgene plasmid #26868) was a gift from Nicholas Gaiano^56^.

### In utero electroporation

*In utero* electroporations were performed as described previously^91^. Survival surgeries were performed on timed pregnant mice (E15.5), to expose their uterine horns. Approximately 1 μL of endotoxin-free plasmid DNA was injected into each embryo’s lateral ventricles. Co-electroporations were performed with the following concentrations for each injection solution: SMOA1-IRES-GFP, DN-RBPJ-IRES-mTagBFP2, NICD-IRES-mTagBFP2, GFP control, BFP control, GliBS-GFP, or Hes5-dsRed at 0.75 mg/mL and CMV-mPB at 0.3 mg/mL. For E15.5 electroporations, five pulses of 100 ms each separated by 950 ms were applied at 45 V. Embryos were put back into the abdominal cavity and the pregnant dams were sutured. Embryos were allowed to develop *in utero* for the indicated time before their brains were dissected.

### Embryonic forebrain slice culture

Whole brains from E15.5 wildtype CD-1 or B6 mice, or NICD mice, were dissected and placed in ice-cold Complete HBSS (1x HBSS, 2.5 mM Hepes, 30 mM D-Glucose, 1 mM CaCl_2_, 1 mM MgSO_4_, 4 mM NaHCO3). Brains were embedded in 3-4% Low Melting Point Agarose dissolved in Complete HBSS and allowed to solidify on ice. Embedded brains were sliced using a vibratome (Leica VT1200S) into 300 μm thick slices and placed in Complete HBSS. Slices were transferred into uncoated Millicell cell culture membrane inserts in 6-well plates and cultured in Slice Culture Media (Complete HBSS, Basal Medium Eagle, 20 mM D-glucose, 1 mM L-glutamine, penicillin-streptomycin) at 37º C, 5% CO2, and 100% humidity. For drug treatments, 100% DMSO was added to the Slice Culture Media for a final concentration of 0.1% DMSO. 100% DMSO was used to make a 10 mM DAPT stock solution. 100% DMSO was also used to make a 100 μg/mL SHH ligand stock solution, which was added to the Slice Culture Media for a final concentration of 100 ng/mL SHH and 0.1% DMSO. For SHH+DAPT treatments, SHH and DAPT stock solutions were added to Slice Culture Media for a final concentration of 10 μM DAPT, 100 ng/mL SHH, and 0.1% DMSO. Slices were plated immediately with 1.5 mL of culture media containing DMSO, SHH only, or SHH+DAPT. After 1 day of incubation, s0.5 mL of culture media was removed and replaced with 0.5 mL of fresh media, using the same drug concentrations. After 2 days *in vitro* (DIV), cell culture media were aspirated, and slices were washed in 1x PBS and fixed in cold 4% PFA for 30 min. Fixed slices were washed twice with 1x PBS and then used for immunohistochemical analysis as described below.

### Immunohistochemistry

Whole embryonic brains were fixed in 4% paraformaldehyde (PFA) for 1 hour at room temperature (RT). Brains were sectioned coronally at 100 μm with a vibrating microtome. Free-floating sections were placed in 24-well plates and blocked with 500 μL of 10% donkey serum and 0.2% Triton-X in 1x PBS for 2 hours at RT. Blocking solution was then removed and sections were incubated with primary antibodies in 10% donkey serum and 0.2% Triton-X in 1x PBS overnight (16 hours) at 4°C. Primary antibody solution was then removed and sections were washed at RT with 1x PBS three times for 5 minutes each. After washing, sections were incubated with secondary antibodies in 1x PBS for 1 hour at RT. Sections were then washed again using 1x PBS three times for 5 minutes each. Sections were mounted on glass slides with ProLong Diamond Antifade Mountant. Images were captured using a LSM900 Zeiss laser scanning confocal microscope in Airyscan 2 Multiplex 4Y mode at 20x or 40x magnification. Antibodies used for immunostaining are listed in the Key Resources Table (**Table S7**). The concentration of each primary antibody used was: goat anti-OLIG2 (1:1000), rat anti-PDGFRA (1:1000), rabbit anti-TagRFP to detect mTagBFP2 (1:500), chicken anti-GFP (1:500). Donkey secondary antibodies conjugated to Alexa Fluor 488, Rhodamine Red-X, Alexa Fluor 647, or Alexa Fluor 405 were used at 1:500.

### Microdissection, tissue dissociation, and MACS

Whole brains from either E16.5 or E17.5 *Emx1-Cre(+/cre);R26-LSL-NICD(+/fl)* or *Emx1-Cre(+/cre)* mouse embryos were collected and microdissected for dorsal forebrain tissue in 1x Neurobasal-A medium under a microscope. Microdissected tissue from 2-5 brains were pooled for each sample and then dissociated with the Worthington Papain Dissociation System according to the Dissociation of Mouse Embryonic Neural Tissue protocol (document CG00053-Rev C) by 10x Genomics. Dissociated samples were strained with 30 μm MACS SmartStrainers into 1x HBSS and 10% FBS solution and centrifuged at 1000 rpm for 5 minutes to pellet the dissociated single cells. Neural progenitor cells were collected for each sample by magnetic-activated cell sorting (MACS) using Anti-Prominin1 microbeads according to the manufacturer’s recommended protocol.

### RNA-seq and analysis

Following MACSorting of dorsal forebrain progenitors, RNA was extracted from each sample of 2-5 pooled brains with the Zymo Quick-RNA microprep kit, following manufacturer’s instructions. RNA concentration was measured with QuBit RNA HS assay kit and submitted for cDNA library preparation and sequencing by the Genomics Shared Resource Facility at the University of Colorado Anschutz Medical Campus (RRID: SCR_021984). Libraries were prepared using SMARTer Stranded Total RNA-seq Kit for low input ribo-depleted RNA and sequenced at 80 million reads/sample using the Illumina NovaSeq X platform. 3 biological replicates were obtained for each of CTRL and NICD E16.5 and E17.5 conditions.

The nf-core RNA-seq pipeline (v 3.14.0)^77^ was used for data preprocessing and analysis with default parameters. STAR^92^ was specified in the nf-core pipeline for alignment to the GRCm38 mouse genome, and Salmon was specified for transcript quantification based on STAR alignments^93^. Gene-level count tables resulting from Salmon were used as input to perform differential gene expression analysis using the DESeq2 R Bioconductor package (v 1.42.1)^78^. Genes were considered differentially expressed if found to have a Benjamini-Hochberg adjusted p-value < 0.05. Exploratory data analysis and principal component analysis were performed with DESeq2 and sheatmaps were generated using the pheatmap package (v 1.012)^79^. GO term enrichment analysis was performed using clusterProfiler package (v 4.10.1)^80^.

### ATAC-seq and analysis

Following MACSorting of dorsal forebrain progenitors, 50,000 cells from each sample were used to prepare an ATAC-seq library using the Zymo-Seq ATAC library kit, following manufacturer’s instruction. Purified DNA library concentrations were measured using the Qubit DNA HS assay kit and submitted to the Genomics Shared Resource Facility at the University of Colorado Anschutz Medical Campus (RRID: SCR_021984) for sequencing with the Illumina NovaSeq X platform at 30 million reads/sample. 3 biological replicates were obtained for CTRL and NICD E16.5 conditions.

The nf-core ATAC-seq pipeline (v 2.1.2)^77^ was used for data preprocessing and alignment to the GRCm38 mouse genome with default parameters. Reads mapped to the mitochondrial genome were removed. Bam files resulting from BWA alignment^94^ in the nf-core pipeline were then filtered for fragments less than 100 bp in length using sambamba (v 0.8.2)^81^, to isolate fragments likely originating from nucleosome-free regions^95^. Newly filtered bam files were then indexed using SAMtools (v 1.16.1)^82^ and used to generate bigwig files with Deeptools (v 3.3.0)^83^ for visualization of accessible chromatin peaks in Integrative Genomics Viewer (IGV)^96^. Peaks were called using MACS2 (2.1.1.20160309)^84^ with the following parameters: -f BAMPE, --keep-dup all, -g mm, call-summits. Principal component analysis was performed and profile plots were generated using Deeptools. Differential chromatin accessibility analysis was carried out using the Diffbind R Bioconductor package (v 3.12.0) and specifying DESeq2 for the analysis^85^. Peaks with FDR < 0.05 were considered significant and differentially accessible. Significant peaks were annotated and assigned to the nearest TSS (−3000 bp, +3000 bp) using the ChIPseeker package (v 1.38.0)^86^. Normalized read count plots were made by extracting the normalized read counts from the Diffbind object, filtering for significant peaks, and assigning peaks to the nearest TSS. Following annotation, peaks were categorized based on curated gene lists (progenitor genes, neurogenic genes, etc) and the average normalized read counts across replicates were plotted. Normality of the data was tested using Shapiro-Wilk’s test, and variance was tested with Levene’s test. For comparisons where the data is not normally distributed, Wilcoxon rank-sum test was used to determine significance. To identify transcription factor binding motifs within differentially accessible sites, we used HOMER (v 4.9)^87^ to perform *de novo* motif enrichment analysis. To locate instances of motifs, we used HOMER findMotifsGenome.pl find, using motif files generated from HOMER *de novo* motif enrichment and the RBPJ motif file from the HOMER. For figures featuring genome tracks, bigwig files for each condition were merged with the “samtools merge” command to visualize averaged peaks.

### Quantification and statistical analysis

All images were captured using a LSM900 Zeiss laser scanning confocal microscope in Airyscan 2 Multiplex 4Y mode at 20x or 40x magnification. All images were exported in tiff or jpeg format with no compression. Brightness, contrast, and background were adjusted equally for the entire image between controls and mutants using the “Brightness/Contrast” and “Levels” function from “Image/Adjustment” options in Adobe Photoshop or Fiji without any further modification.

For all immunostainings, 3 or more histological sections at three distinct rostral-caudal z-planes from each of 3-5 different animals (at least nine sections total for each condition) were analyzed in the entire dorsal pallium. Biological replicates are individual animals. For slice cultures, biological replicates were individual slices from 3-4 different animals that were histologically matched at the rostral-caudal level between untreated and treated groups. Confocal, single-plane optical sections were used for quantification. Cells were analyzed in columns spanning the entire dorsal pallium across the lateral-medial axis. Total number of marker-positive cells were quantified from these columns in Fiji/ImageJ^76^ and then divided by the area of a column to get cell density (cells/mm^2^). The cell density was averaged across multiple columns and sections for each animal. For CTRL and NICD E17.5 brains, the cell density was quantified within the ventricular and subventricular zones. For electroporations, the percentage of electroporated cells that were positive for a marker was calculated and averaged for each animal. All statistical tests were performed using R. Normality of the data was tested using Shapiro-Wilk’s test, and variance was tested with Levene’s test. For independent two-group experiments, an unpaired two-tailed Student’s t-test was used to determine statistical significance between two groups with equal variance. For comparisons between two groups with unequal variance, Welch’s t-test was used. For analysis involving 3 or more independent groups with equal variance, a one-way ANOVA was used followed by Tukey’s *post hoc* test. For analysis involving 3 or more independent groups with unequal variance, a Welch ANOVA test was used followed by Games-Howell’s *post hoc* test. For analysis involving 3 or more independent groups that were not normally distributed, a Kruskal-Wallis test was used followed by Dunn’s *post hoc* test. Values were considered statistically significant at p < 0.05.

For reporter assay experiments, images were taken with the same settings across all sections from all conditions. 3 distinct z-planes from 2 or more histological sections at matching rostral-caudal areas from each of 3 different animals were analyzed. Images were exported in tiff format with no compression and with all 3 channels (RGB). For each image, a box of width 1000 px and height 210 px (approximately 100 μm) was drawn over the ventricular and subventricular zones in the dorsal pallium, with the ventricular wall at bottom of the box. Based on this box, the image was cropped and saved as a new tiff image. Images in figures were linearly adjusted with increased brightness for the purpose of visibility in the figure. Analysis was performed on images without any adjustments. The fluorescence intensity of each image was analyzed using Fiji/ImageJ by drawing a Region of Interest (ROI) around every visible cell in the ventricular and subventricular zone in the cropped image; measuring the mean fluorescence intensity values for the red, green, and blue channels for each ROI; and saving the values in an excel spreadsheet file. Values for the red and green channels were each normalized to the corresponding values for the blue channel before statistical testing was performed using R. Biological replicates were individual cells circled as ROIs from 3 different animals. Normality was tested using Shapiro-Wilk’s test, and the equality of variances were tested using Levene’s test. For data that did not meet the assumptions for one-way ANOVA, the Kruskal-Wallis test was used followed by Dunn’s *post hoc* test to determine significance between conditions. P-values were adjusted using the Bonferroni method. Values were considered statistically significant at p < 0.05. To determine the correlation of red and green values (normalized to blue), Pearson’s correlation coefficient was calculated.

## RESOURCE AVAILABILITY

### Lead Contact

Further information and requests for resources and reagents should be directed to and will be fulfilled by the lead contact, Santos Franco. All reagents developed in this study can be obtained with a completed material transfer agreement.

## ACKNOWLEDGEMENTS

This work was supported by NIH/NINDS R01 NS124166 (S.J.F., B.H.A., C.G.S.), NIH/NINDS R01 NS109239 (S.J.F.), NIH/NINDS F31 NS129289 (L.N.T.), NIH/NIGMS T32 GM136444 (L.N.T.), NIH/NIGMS T32 GM136444-S1 (L.N.T.), and the Gates Summer Internship Program through the Gates Institute (A.Sa.). We thank Rebecca O’Rourke and Dr. Tyler Gibson for technical advice for bioinformatics analyses. We thank the following people for helpful scientific discussions and critical reading of the manuscript: Dr. Joseph Brzezinski, Dr. Adam Almeida, Dr. Kimberly Arena, Madisen Mason, Aleezah Balolia, J.P. Martin, Salvador Guerra, and students of the Cell & Developmental Biology Advanced Writing Workshop.

## AUTHOR CONTRIBUTIONS

Conceptualization, L.N.T., S.J.F., B.H.A., C.G.S.; Investigation, L.N.T., A.Sh., K.H.S., A.Sa., E.D., M.J.W., T.J.I., S.A.N., S.J.F.; Formal Analysis, L.N.T.; Resources, F.G.-M.; Writing Original Draft, L.N.T., S.J.F.; Visualization, L.N.T.; Funding Acquisition, L.N.T, S.J.F., B.H.A., C.G.S.

## SUPPLEMENTAL INFORMATION

Document S1. Figures S1-S2, Table S2, and Table S7.

Table S1. Excel file containing E16.5 RNA-seq data too large to fit into a PDF. Related to Figure 2.

Table S3. Excel file containing ATAC-seq data too large to fit into a PDF. Related to Figure 3.

Table S4. Excel file containing ATAC-seq data related to neurogenic transcription factor binding motifs. Related to Figure 4.

Table S5. Excel file containing ATAC-seq data related to gliogenic transcription factor binding motifs. Related to Figure 4.

Table S6. Excel file containing E17.5 RNA-seq data too large to fit into a PDF. Related to Figure 5.

