## Supplemental Document S1 for "Epigenetic priming of neural progenitors by Notch enhances Sonic hedgehog signaling and establishes gliogenic competence"

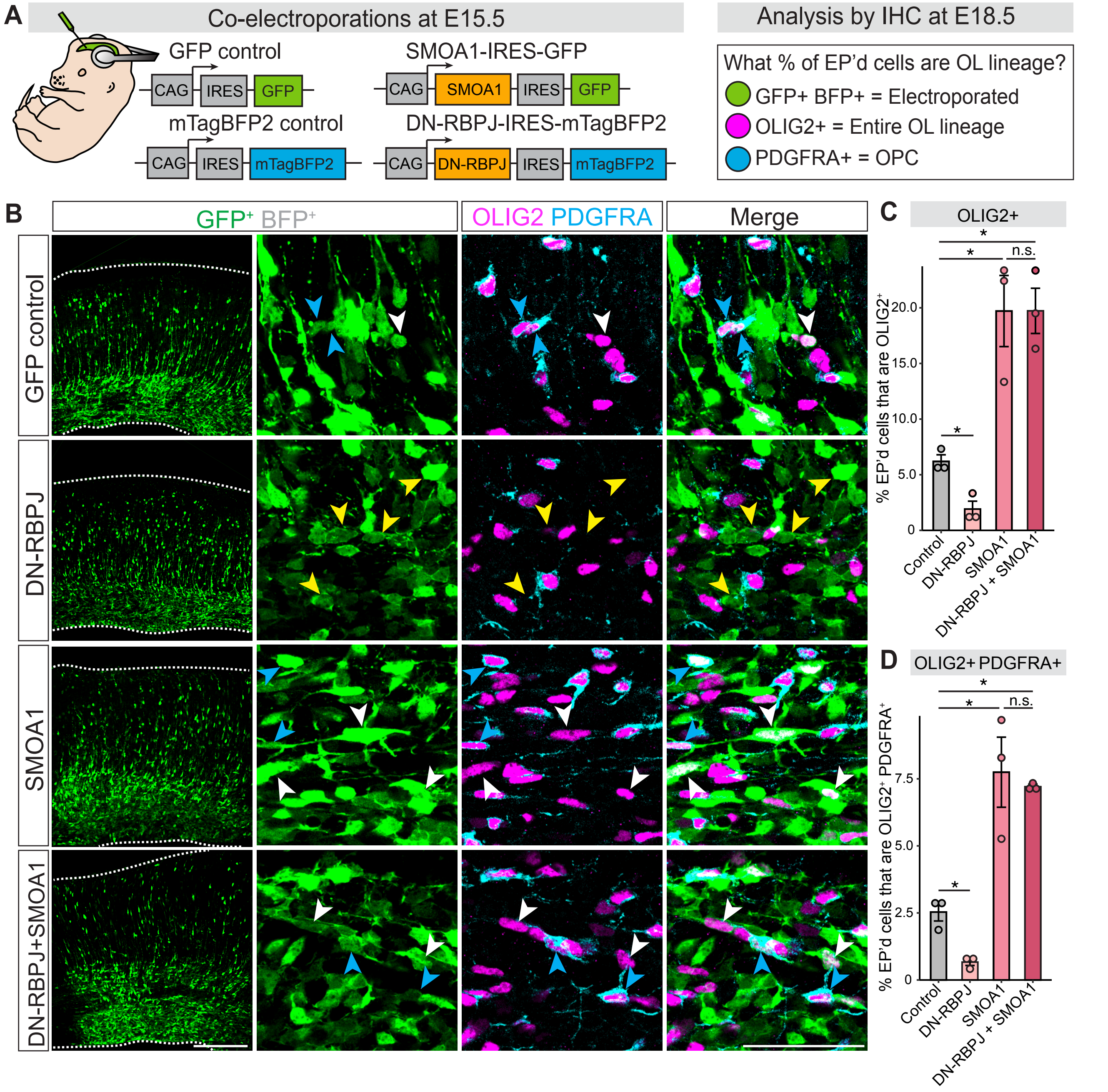


**Supplemental Figure 1 (related to Figure 1). The requirement for Notch signaling in oligodendrocyte lineage cell production is bypassed by increasing SHH signaling.** (A) Schematic of in utero electroporation approach. Wildtype mouse embryos were co-electroporated with GFP + BFP control, DN-RBPJ-IRES-mTagBFP2 + GFP, SMOA1-IRES-GFP + BFP, or DN-RBPJ + SMOA1 at E15.5. At E18.5, brains were dissected and analyzed by IHC for OLIG2 and PDGFRA to identify oligodendrocyte lineage cells. OL, oligodendrocyte; OPC, oligodendrocyte precursor cell. EP'd = electroporated. IHC = immunohistochemistry. (B) Left panels show overview images of the electroporations in the dorsal pallium. Since electroporated cells are both GFP+ and BFP+, the BFP channel is not shown. Dotted lines outline the dorsal (top) and ventral (bottom) limits of the pallium. Scale bar, 200 m. Representative higher magnification images of brains electroporated with GFP control, DN-RBPJ, SMOA1, and SMOA1+DN-RBPJ. White arrowheads denote GFP+ OLIG2+ cells, blue arrowheads denote GFP+ OLIG2+ PDGFRA+ cells, and yellow arrowheads denote GFP+ OLIG2- PDGFRA- cells. Scale bar, 50 m (C-D) Quantification of oligodendrocyte lineage cells among electroporated cells. Graphs show the average percentage (± SEM among biological replicates) of electroporated (GFP+ BFP+) cells that were OLIG2+ (C) or OLIG2+ PDGFRA+ (D). For comparisons between Control and DN-RBPJ, either Student's t-test for equal variance or Welch's t-test for unequal variance was performed. Welch's t-test Control vs DN-RBPJ: (C) *p= 0.009; Student's t-test Control vs DN-RBPJ (D) *p= 0.0063. For comparisons between Control, SMOA1, and DN-RBPJ+SMOA1, one-way ANOVA and Tukey's post-hoc tests were performed. One-way ANOVA (C) p = 0.0073; (D) p = 0.006. Tukey's post-hoc test: *p< 0.05, n.s. = not significant. N= 3 brains for each condition.

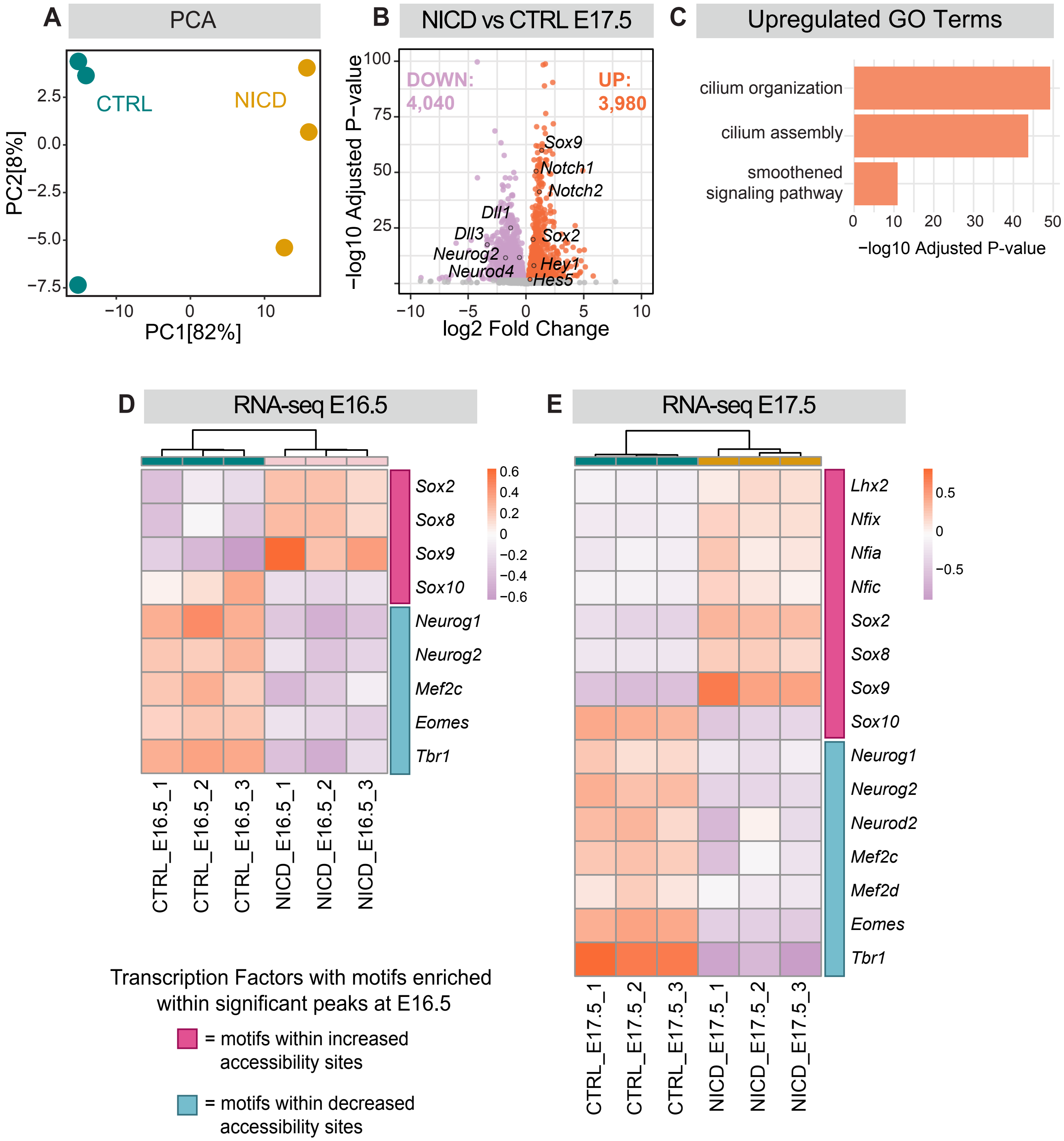


**Supplemental Figure 2 (related to Figure 5). RNA-seq on dorsal progenitors at E17.5 reveals upregulation of cilia and Smoothened pathway genes, as well as transcription factors whose motifs were enriched in differentially accessible chromatin regions.** (A) PCA plot indicating separation of CTRL (N= 3) and NICD (N= 3) E17.5 transcriptomes by condition. PCA = Principal component analysis. (B) Volcano plot representing differentially expressed genes between CTRL and NICD. Differentially expressed genes with adjusted p-value < 0.05 were considered to be significant. 4,040 genes were significantly downregulated (purple) and 3,980 genes were upregulated (orange) in NICD compared to CTRL. Selected Notch-related genes are outlined and labeled in the plot. (C) GO term enrichment analysis of upregulated genes. X-axis shows the -log Adjusted P-value, with the y-axis showing selected GO terms. GO = gene ontology. (D-E) Heatmaps representing selected DEGs at E16.5 (D) or E17.5 (E) for transcription factors whose binding motifs were enriched in either increased or decreased accessibility sites found in E16.5 ATAC-seq analysis. Heatmap scales show variance-stabilized gene expression levels, with colors representing relative expression levels across samples. DEG = differentially expressed genes.

**Table S2. RNA-seq results of Shh pathway genes.**

| ensembl | symbol | entrez | baseMean | log2FoldChange | lfcSE | stat | pvalue | padj |
|---|---|---|---|---|---|---|---|---|
| ENSMUSG00000025407 | Gli1 | 14632 | 581.078868 | -1.8778382 | 0.16737024 | -11.219667 | 3.26E-29 | 1.88E-26 |
| ENSMUSG00000028681 | Ptch2 | 19207 | 94.6735435 | -2.7297505 | 0.41401621 | -6.5933421 | 4.30E-11 | 3.70E-09 |
| ENSMUSG00000064325 | Hhip | 15245 | 63.2238949 | -1.1883386 | 0.43180108 | -2.752051 | 0.00592233 | 0.05630482 |
| ENSMUSG00000022687 | Boc | 117606 | 5761.54333 | 0.31415163 | 0.11931719 | 2.63291164 | 0.00846564 | 0.07222784 |
| ENSMUSG00000021318 | Gli3 | 14634 | 11172.1643 | 0.31288922 | 0.12787084 | 2.44691622 | 0.01440843 | 0.10451448 |
| ENSMUSG00000021466 | Ptch1 | 19206 | 1373.28588 | -0.4604268 | 0.19052273 | -2.41665 | 0.01566407 | 0.11051796 |
| ENSMUSG00000038119 | Cdon | 57810 | 17770.6904 | 0.26583257 | 0.11009338 | 2.41461001 | 0.01575207 | 0.11092601 |
| ENSMUSG00000073791 | Efcab7 | 230500 | 132.879962 | -0.5873952 | 0.27680881 | -2.1220249 | 0.03383565 | 0.18498893 |
| ENSMUSG00000001761 | Smo | 319757 | 3615.28976 | 0.23384972 | 0.1174593 | 1.99090005 | 0.04649188 | 0.22612738 |
| ENSMUSG00000022812 | Gsk3b | 56637 | 13348.019 | 0.226223 | 0.11723617 | 1.92963491 | 0.05365209 | 0.24740702 |
| ENSMUSG00000025231 | Sufu | 24069 | 3151.59319 | -0.1910889 | 0.11960295 | -1.5976942 | 0.11011105 | 0.38264263 |
| ENSMUSG00000048402 | Gli2 | 14633 | 4417.92676 | 0.15682448 | 0.10582652 | 1.48190146 | 0.13836652 | 0.43044689 |
| ENSMUSG00000004364 | Cul3 | 26554 | 4830.74045 | 0.13286624 | 0.0969723 | 1.37014631 | 0.17064123 | 0.47997396 |
| ENSMUSG00000050382 | Kif7 | 16576 | 2081.50887 | 0.1225576 | 0.11605069 | 1.05606957 | 0.29093643 | 0.62120497 |
| ENSMUSG00000030768 | Disp1 | 68897 | 581.726667 | -0.1391591 | 0.17032877 | -0.8170029 | 0.41392678 | 0.72566349 |
| ENSMUSG00000052957 | Gas1 | 14451 | 2892.16603 | -0.0217583 | 0.19563264 | -0.1112203 | 0.91144165 | 0.97441998 |
| ENSMUSG00000036555 | Iqce | 74239 | 2635.17641 | 0.0041126 | 0.11389279 | 0.0361094 | 0.97119513 | 0.98973221 |

**Table S7. Key resources table.**

| REAGENT or RESOURCE | SOURCE | IDENTIFIER |
|---|---|---|
| Antibodies | | |
| Goat anti-OLIG2 | R&D Systems | Cat# AF2418; RRID: AB_2157554 |
| Rat anti-PDGFRA | Thermo Fisher Scientific | Cat# # **720219;** RRID: AB_2633205 |
| Rabbit anti-TagRFP | Thermo Fisher Scientific | Cat# R10367; RRID: AB_10563941 |
| Chicken anti-GFP | Thermo Fisher Scientific | Cat# A10262; RRID: AB_2534023 |
| Donkey secondary antibody anti-Chicken Alexa Fluor 488 | Jackson ImmunoResearch | Cat# 703-545-155; RRID: AB_2340375 |
| Donkey secondary antibody anti-Goat Alexa Fluor 647 | Jackson ImmunoResearch | Cat# 705-605-147; RRID: AB_2340437 |
| Donkey secondary antibody anti-Rabbit Alexa Fluor 405 | Jackson ImmunoResearch | Cat# 711-475-152; RRID: AB_2340616 |
| Donkey secondary antibody anti-Rat Rhodamine Red-X | Jackson ImmunoResearch | Cat# 712-297-003; RRID: AB_2340679 |
| DAPI | Thermo Fisher Scientific | Cat# D1306 |
| Chemicals, peptides, and recombinant proteins | | |
| DMSO | Millipore Sigma | Cat# D2650-100ML |
| DAPT | Millipore Sigma | Cat# 565770-5MG |
| SHH ligand (C25II), mouse | Genscript | Cat# Z03050 |
| 10x HBSS | Thermo Fisher Scientific | Cat# 14-185-052 |
| 1 M Hepes | Thermo Fisher Scientific | Cat# 15-630-106 |
| 1 M D-Glucose | Thermo Fisher Scientific | Cat# J60067 |
| CaCl2 | Millipore Sigma | Cat# C4901 |
| MgSO4 | Millipore Sigma | Cat# M2643-500G |
| NaHCO3 | Millipore Sigma | Cat# S5761-500G |
| Basal Medium Eagle | Millipore Sigma | Cat# B1522-500ML |
| 200 mM L-glutamine | Gemini Bio | Cat# 400-106 |
| Penicillin-Streptomycin | Lonza Bioscience | Cat# DE17-602E |
| Millicell cell culture plate inserts | Millipore Sigma | Cat# Z353086 |
| Cell culture 6-well plates | Greiner Bio-one | Cat# 657160 |
| Neurobasal-A Medium (1X) | Gibco | Cat# 12349-015 |
| Critical commercial assays | | |
| NEBuilder HiFi DNA Assembly Master Mix | New England Biolabs | Cat# E2621S |
| ProLong Diamond Antifade Mountant | Thermo Fisher Scientific | Cat# P36961 |
| Monarch MiniPrep Kit | New England Biolabs | Cat# T1010L |

| | | |
|---|---|---|
| EndoFree Plasmid Maxi Kit | Qiagen | Cat# 12362 |
| Papain Dissociation System | Worthington Biochem | Cat# LK003150 |
| MACS SmartStrainers (30 μm) | Miltenyi Biotec | Cat# 130-098-458 |
| Magnetic Separation columns | Miltenyi Biotec | Cat# 130-042-201 |
| MACS MultiStand | Miltenyi Biotec | Cat# 130-042-303 |
| Anti-Prominin-1 MicroBeads mouse | Miltenyi Biotec | Cat# 130-092-333 |
| Zymo-seq ATAC library kit | Zymo | Cat# D5458 |
| Quick-RNA Microprep kit | Zymo | Cat# R1050 |
| Genomics Shared Resource | University of Colorado | RRID: SCR_021984 |
| Deposited data | | |
| Raw RNA-seq fastq files | This paper | To be deposited on GEO repository database |
| Raw ATAC-seq fastq files | This paper | To be deposited on GEO repository database |
| Experimental models: Organisms/strains | | |
| B6 (C57BL/6J) | The Jackson Laboratory | stock no. 000664 |
| R26-LSL-NICD (*Gt(ROSA)26Sor*$^{tm1(Notch1)Dam}$/J (ROSA26$^{loxP\text{-}stop\text{-}loxP\text{-}Notch1\text{-}ICD}$) | The Jackson Laboratory | stock no. 008159 |
| Emx1-Cre (B6.129S2-Emx1tm1(cre)Kri/J) | The Jackson Laboratory | stock no. 005628 |
| Crl:CD1(ICR**)** | Charles River | strain no. 022 |
| Recombinant DNA | | |
| CMV-mPB | Winkler et al., 2018 | N/A |
| pPB-CAG-IRES-GFP | Tran et al., 2023 | N/A |
| pPB-DN-RBPJ-IRES-mTagBFP2 | Tran et al., 2023 | N/A |
| pPB-NICD-IRES-mTagBFP2 | This paper | N/A |
| pPB-SMOA1-IRES-GFP | This paper | N/A |
| pPB-CAG-IRES-mTagBFP2 | This paper | N/A |
| pGL3b-8xGliBS:EGFP | Hyman et al.[55] | Cat# 84602; RRID: Addgene_84602 |
| Hes5p-dsRed | Mizutani et al.[56] | Cat# 26868; RRID: Addgene_26868 |
| Software and algorithms | | |
| Fiji/ImageJ | Schneider et al.[76] | https://fiji.sc |
| Photoshop | Adobe | Adobe.com |
| Illustrator | Adobe | Adobe.com |
| R Studio | Posit | https://posit.co/download/rstudio-desktop/ |
| Integrative Genomics Viewer (IGV) | IGV | https://igv.org/doc/desktop/ |
| nf-core RNA-seq pipeline | Ewels et al.[77] | https://github.com/nf-core/rnaseq |
| nf-core ATAC-seq pipeline | Ewels et al.[77] | https://github.com/nf-core/atacseq |
| DESeq2 | Love et al.[78] | https://github.com/theloveland/DESeq2 |

| | | |
|---|---|---|
| pheatmap | Kolde et al.[79] | https://github.com/raivokolde/pheatmap |
| clusterProfiler | Yu et al.[80] | https://github.com/YuLab-SMU/clusterProfiler |
| Sambamba | Tarasov et al.[81] | https://github.com/biod/sambamba |
| SAMtools | Li et al.[82] | https://github.com/samtools |
| Deeptools | Ramírez et al.[83] | https://github.com/deeptools/deepTools |
| MACS2 | Zhang et al.[84] | https://github.com/macs3-project/MACS |
| Diffbind | Stark et al.[85] | https://github.com/hnthirima/DiffBind |
| ChIPseeker | Yu et al.[86] | https://github.com/YuLab-SMU/ChIPseeker |
| HOMER | Heinz et al.[87] | https://github.com/javrodriguez/HOMER |
